# Direct Tests of Cytochrome Function in the Electron Transport Chain of Malaria Parasites

**DOI:** 10.1101/2023.01.23.525242

**Authors:** Tanya J. Espino-Sanchez, Henry Wienkers, Rebecca G. Marvin, Shai-anne Nalder, Aldo E. García-Guerrero, Peter E. VanNatta, Yasaman Jami-Alahmadi, Amanda Mixon Blackwell, Frank G. Whitby, James A. Wohlschlegel, Matthew T. Kieber-Emmons, Christopher P. Hill, Paul A. Sigala

## Abstract

The mitochondrial electron transport chain (ETC) of *Plasmodium* malaria parasites is a major antimalarial drug target, but critical cytochrome functions remain unstudied and enigmatic. Parasites express two distinct cyt *c* homologs (*c* and *c*-2) with unusually sparse sequence identity and uncertain fitness contributions. *P. falciparum* cyt *c*-2 is the most divergent eukaryotic cyt *c* homolog currently known and has sequence features predicted to be incompatible with canonical ETC function. We tagged both cyt *c* homologs and the related cyt *c*_1_ for inducible knockdown. Translational repression of cyt *c* and cyt *c*_1_ was lethal to parasites, which died from ETC dysfunction and impaired ubiquinone recycling. In contrast, cyt *c*-2 knockdown or knock-out had little impact on blood-stage growth, indicating that parasites rely fully on the more conserved cyt *c* for ETC function. Biochemical and structural studies revealed that both cyt *c* and *c*-2 are hemylated by holocytochrome *c* synthase, but UV-vis absorbance and EPR spectra strongly suggest that cyt *c*-2 has an unusually open active site in which heme is stably coordinated by only a single axial amino-acid ligand and can bind exogenous small molecules. These studies provide a direct dissection of cytochrome functions in the ETC of malaria parasites and identify a highly divergent *Plasmodium* cytochrome *c* with molecular adaptations that defy a conserved role in eukaryotic evolution.

**SIGNIFICANCE STATEMENT:** Mitochondria are critical organelles in eukaryotic cells that drive oxidative metabolism. The mitochondrion of *Plasmodium* malaria parasites is a major drug target that has many differences from human cells and remains poorly studied. One key difference from humans is that malaria parasites express two cytochrome *c* proteins that differ significantly from each other and play untested and uncertain roles in the mitochondrial electron transport chain (ETC). Our study revealed that one cyt *c* is essential for ETC function and parasite viability while the second, more divergent protein has unusual structural and biochemical properties and is not required for growth of blood-stage parasites. This work elucidates key biochemical properties and evolutionary differences in the mitochondrial ETC of malaria parasites.

## INTRODUCTION

Malaria is a devastating human disease that remains a pressing global health challenge. Symptoms of malaria are caused by red blood cell (RBC) infection by eukaryotic single-celled parasites of the genus *Plasmodium*, with most severe infections and deaths due to *Plasmodium falciparum*. RBCs are the most heme-rich cell in the human body, and heme metabolism is central to blood-stage parasite biology and therefore a potential vulnerability (1–3). Indeed, drugs that interfere or interact with parasite heme metabolism have historically provided potent antimalarial therapies, including chloroquine, atovaquone, and artemisinin (4). During their 48-hour life cycle after RBC invasion, malaria parasites internalize and proteolytically degrade up to 80% of red cell hemoglobin (5). This massive catabolic process liberates prodigious amounts of labile heme that is detoxified via biomineralization into inert crystalline hemozoin within the acidic parasite digestive vacuole (Fig. 1). 4-Aminoquinoline drugs such as chloroquine have potent antimalarial activity via their consensus mechanism of blocking hemozoin formation that results in accumulation of toxic labile heme (4, 6).

**Figure 1.**
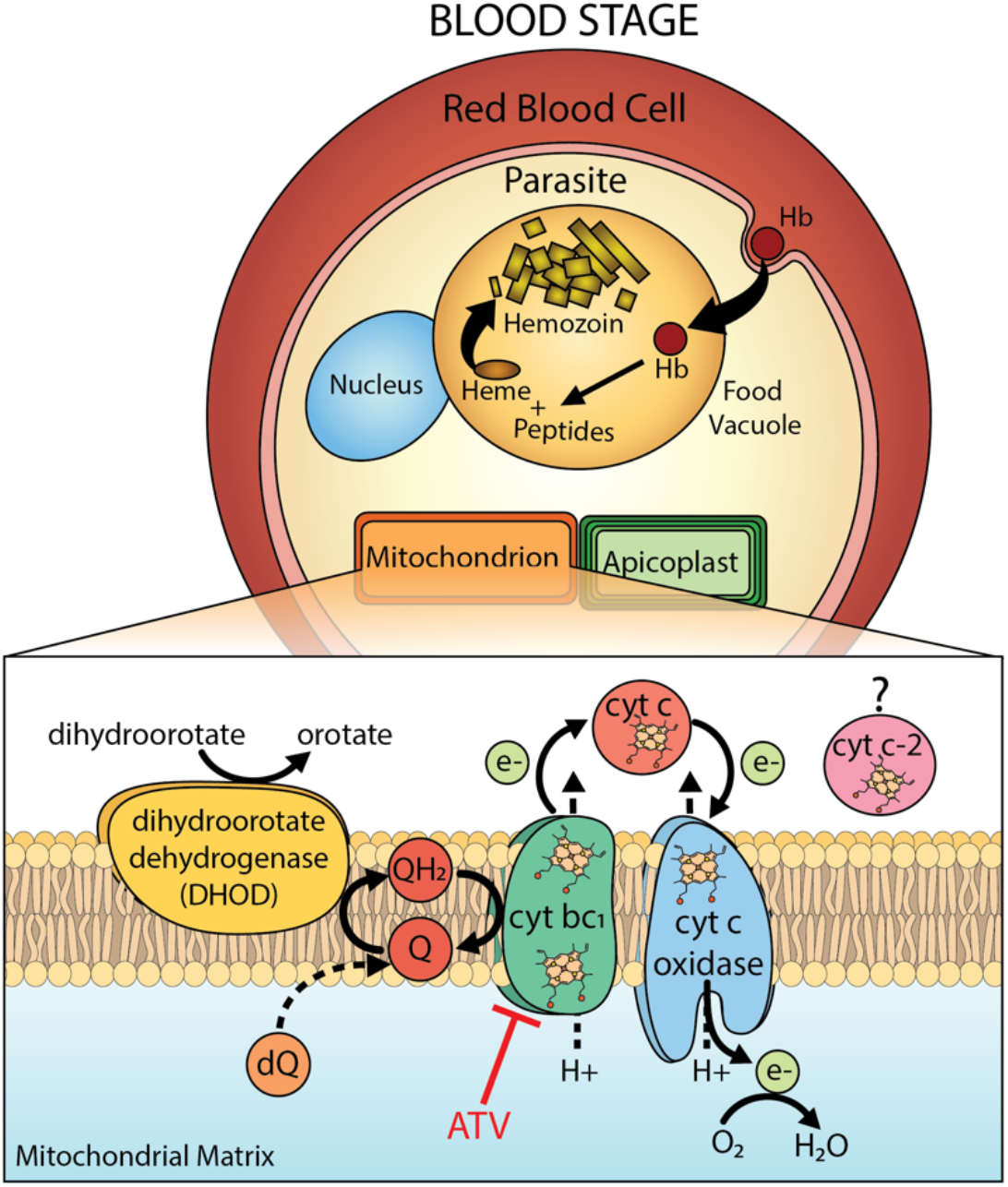
Schematic model of heme metabolism during blood-stage infection by a *P. falciparum* parasite, focusing on the critical roles for heme as an electron-transfer co-factor in the mitochondrial respiratory chain. Question mark indictaes uncertainty in the function of the divergent cyt *c*-2 homolog. ATV = atovaquone, Q = ubiquinone, QH_2_ = ubiquinol, dQ = decyl-ubiquinone. Figure adapted from (19).

*P. falciparum* also requires heme as an essential metabolic cofactor, although cellular uses are sparsely studied and surprisingly few heme proteins are annotated in the parasite genome (2, 3). Parasites retain a mitochondrial electron transport chain (ETC) with heme-dependent cytochromes *b* and c_*1*_ in complex III, cytochrome (cyt) *c* as mobile electron carrier between complexes III and IV, and the heme A-containing CoxI subunit of complex IV (Fig. 1). Complex III binds reduced ubiquinol, oxidizes it to ubiquinone, and couples sequential electron transport via cyt *b*, Rieske 2Fe-2S, cyt *c_1_*, cyt *c* and complex IV to proton translocation that polarizes the inner mitochondrial membrane, through a reaction series called the Q cycle (7).

ETC function is essential for parasite viability, and drugs such as atovaquone or endochin-like quinolones (ELQs) that block ubiquinone binding to cyt *b* are lethal to *P. falciparum* (8, 9). However, parasites have redundant mechanisms to polarize the mitochondrial membrane (10), and ATP synthase is dispensable for blood-stage parasites (11), which rely on glycolysis rather than oxidative phosphorylation for ATP synthesis (12, 13). Thus, the sole essential function of the mitochondrial ETC during RBC infection is thought to be oxidative recycling of ubiquinone (10). Indeed, exogenous decyl-ubiquinone (dQ) is sufficient to rescue parasites from the lethal effects of atovaquone (14) or loss of the Rieske Fe-S protein (15). Although multiple mitochondrial dehydrogenases require ubiquinone as an electron acceptor, dihydroorotate dehydrogenase (DHOD) appears to be the most critical of these enzymes since heterologous expression of ubiquinone-independent yeast DHOD rescues *P. falciparum* from atovaquone or ELQ toxicity (9, 10). The special vulnerability of heme-dependent mitochondrial functions for targeting *P. falciparum* in multiple developmental stages was emphasized by a recent high-throughput drug screen that identified mitochondrial ETC and DHOD inhibitors as the largest class of compounds showing dual efficacy against blood- and liver-stage parasites (16).

Heme-dependent cytochromes of complexes III and IV are key components of the ETC, but their functions have not been directly studied. *P. falciparum* also curiously encodes two divergent cyt *c* homologs, and their functions and essentiality for parasite growth remain uncertain. *C*-type cytochromes like cyt *c* and the related cyt *c_1_* covalently bind heme via conserved Cys residues that form stable thioether bonds to the side-chain vinyl groups of heme, in contrast to other proteins that exclusively bind heme via non-covalent interactions. Covalent heme binding depends on the additional mitochondrial protein, holocytochrome *c* synthase (HCCS) (17). Although humans encode a single bi-functional HCCS protein that attaches heme to both cyt *c* and *c_1_*, *Plasmodium* encodes two HCCS homologs thought to be specific for hemylation of cyt *c* (HCCS) or cyt *c_1_* (HCC_1_S) but whose specificities remain untested (2, 18).

To test and distinguish the functional importance of *c*-type cytochromes for *P. falciparum*, we used CRISPR/Cas9 to tag cyt *c_1_* and both cyt *c* homologs for conditional knockdown. These studies reveal that cyt *c_1_* (PF3D7_1462700) and the more conserved cyt *c* homolog (PF3D7_1404100) are essential for ETC function and parasite viability. The divergent cyt *c*-2 (PF3D7_1311700), however, is dispensable for blood-stage parasite growth. Despite their substantial sequence divergence, both cyt *c* homologs are selectively hemylated by the parasite HCCS (PF3D7_1224600) but not HCC_1_S (PF3D7_1203600). Finally, biophysical studies strongly suggest that the divergent cyt *c*-2 binds ferric heme in an unusual penta-rather than hexa-coordinate ligation environment, despite conservation of axial His and Met ligands, and has a binding pocket that exposes heme to exogenous ligands. This work provides a direct functional dissection of mitochondrial cytochromes in malaria parasites, identifies *c*-type cytochrome hemylation as a potential target for antimalarial drug discovery, and uncovers a divergent eukaryotic cytochrome *c* (*c*-2) with highly unusual molecular adaptations and physical properties.

## RESULTS

### Mitochondrial localization of *c*-type cytochromes and holocytochrome c synthases

Because localization data have not previously been reported for the annotated *P. falciparum c*-type cytochromes or holocytochrome *c* synthases, we tested the mitochondrial targeting of these proteins by live-cell microscopy. We created Dd2 *P. falciparum* lines that episomally expressed cyt *c*, cyt *c*-2, cyt *c_1_*, HCCS, or HCC_1_S with a C-terminal GFP tag. For all five proteins, the fluorescence signal observed for GFP co-localized with signal for MitoTracker Red (Fig. 2 and Fig. S1), confirming mitochondrial targeting. This localization is consistent with general biochemical expectations for these proteins and with a recent proteomic study that identified cytochromes *c* and *c_1_* in *P. falciparum* mitochondrial extracts (20). Our studies of endogenously tagged cyt *c* and *c*-2 using immunoprecipitation/mass spectrometry also strongly support mitochondrial localization of these proteins (see below). Live-cell images of parasites expressing cyt *c*_1_-GFP acquired on a higher-resolution Airyscan confocal microscope (Fig. S2) further supported mitochondrial but could not distinguish sub-organellar targeting to the intermembrane space (IMS), the expected localization of *c*-type cytochromes and HCCS homologs.

**Figure 2.**
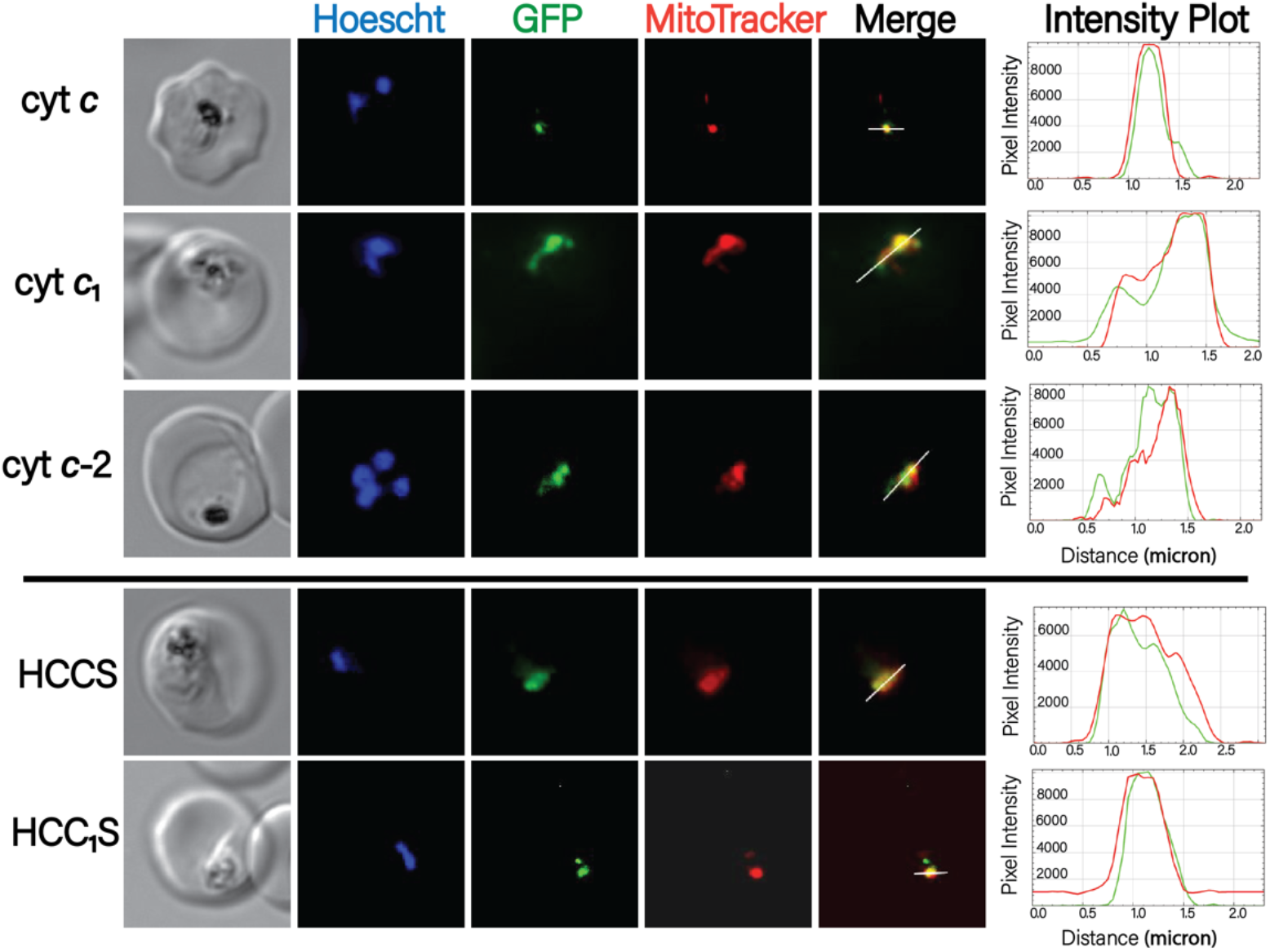
Localization of *c*-type cytochromes and holocytochrome *c* synthase proteins in blood-stage parasites. Images were obtained by live epifluorescence microscopy of parasites expressing episomal second copies of each protein with a C-terminal GFP tag and stained with MitoTracker Red (mitochondrion) and Hoescht (nucleus). The intensity plot to the right of each image series displays the overlap in pixel intensity for GFP and MitoTracker Red as a function of distance along the white line in the merged image.

### Divergent sequence features of *P. falciparum* cytochrome c homologs

We first focused on the two unusually divergent cyt *c* homologs in *P. falciparum*. Cytochrome *c* is one of the most highly conserved proteins in eukaryotes due to its centrality in mitochondrial ETC function and cellular metabolism. Indeed, its sequence variation among distinct organisms has served as a paradigm for understanding protein evolution and as a biomarker for establishing phylogenetic relationships (21–23). To assess if the two annotated *P. falciparum* cyt *c* homologs contain the expected sequence features for conserved function within the mitochondrial ETC, we aligned the sequences of both proteins and human cytochrome *c* (Uniprot P99999).

The primary sequence of *P. falciparum* cyt *c* (PF3D7_1404100) contains a short, parasite-specific N-terminal extension of 11 residues but is otherwise 64% identical to human cyt *c* and retains the expected molecular signatures for a conserved role in mitochondrial electron transport (Fig. 3A). These features include (i) the CXXCH motif comprised of the two Cys residues that form covalent thioether bonds to the vinyl side-chains of heme and the His residue that serves as an axial heme ligand (Fig. 3B), (ii) Lys and Phe residues on a predicted α-helix that is N-terminal to the CXXCH motif and which are critical for HCCS recognition and covalent heme attachment (17, 24), (iii) the C-terminal Met81 residue (human cyt *c* numbering) that serves as the second axial heme ligand (Fig. 3B), and (iv) nearly complete identity of heme-crevice residues immediately N-terminal to Met81 (especially Lys73) that position Met81 for optimal heme coordination and mediate binding to cyt *c_1_* in ETC complex III (7, 21, 25). This high level of sequence conservation suggests that PF3D7_1404100 may retain canonical cyt *c* function as mobile electron carrier in the mitochondrial ETC.

**Figure 3.**
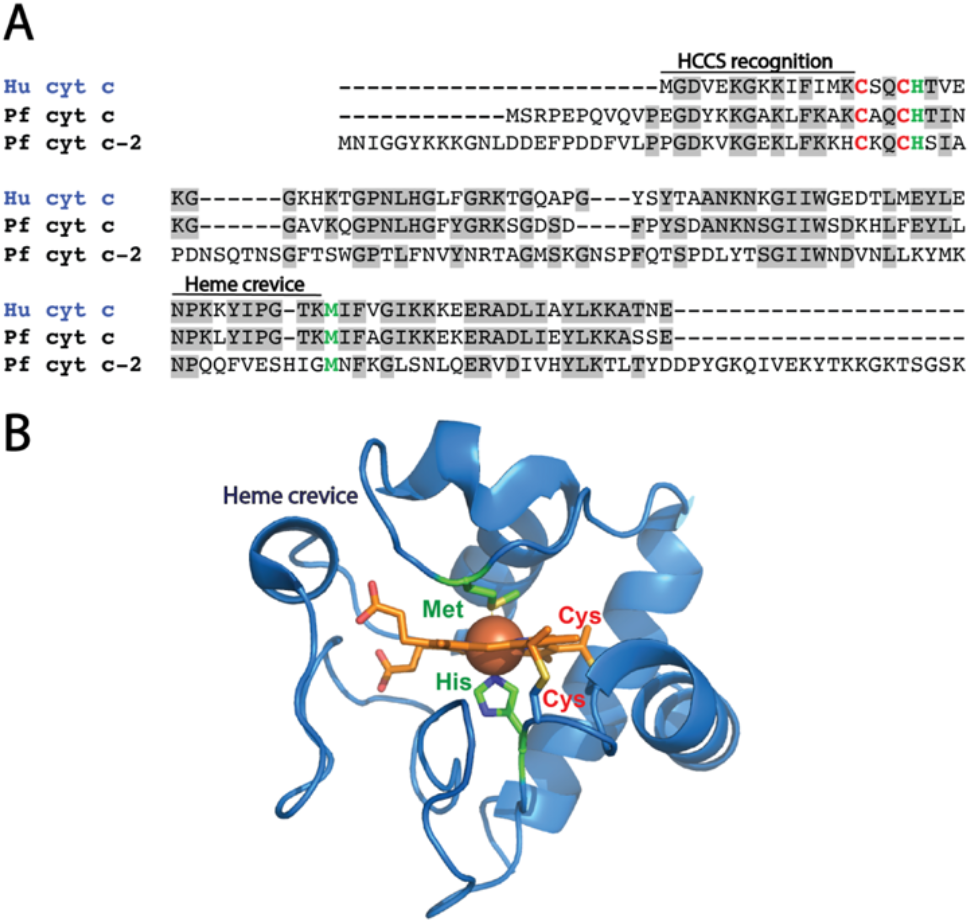
Sequence and structure of cyt *c*. (A) Sequence comparison of human and *Plasmodium* cytochromes *c*. Identical residues are in gray. (B) X-ray crystallographic structure human cyt *c* with bound heme (PDB 3NWV). Cys residues that form thioether bonds to heme are labeled in red. Conserved axial His and Met ligands are in green.

In contrast to *P. falciparum* cyt *c*, the cyt *c*-2 homolog (PF3D7_1311700) shares less than 32% identity with human cyt *c* and features extended, parasite-specific N- and C-termini as well as two short insertions (Fig. 3A) that further reduce overall sequence identity. This low level of sequence conservation for a eukaryotic cyt *c* homolog is highly unusual, and *P. falciparum* cyt *c*-2 is the most divergent eukaryotic cyt *c* homolog currently known (21). Cyt *c*-2 retains the key N-terminal HCCS recognition residues, the CXXCH motif, and Met81, which suggests that this protein is hemylated in the parasite mitochondrion and has the expected biaxial His/Met heme ligands. However, the heme crevice region in cyt *c*-2 has completely diverged from human and parasite cyt *c*, including loss of Lys73 and the curious insertion of a His residue upstream of Met81 (Fig. 3A). Mutations in this region, which is the most highly conserved sequence segment in eukaryotic cyt *c* homologs (21, 22), have been shown to destabilize axial coordination of heme by Met81, perturb the cyt *c* reduction potential, and weaken cyt *c* binding to ETC complex III (25, 26). The total sequence divergence of the heme crevice region in *P. falciparum* cyt *c*-2 strongly suggests that this protein is likely to have an altered heme coordination environment, perturbed reduction potential, and overall physical properties that are not optimized to mediate electron transfer between ETC complexes III and IV. Based on these sequence differences, we predicted cyt *c*-2 to have a divergent function distinct from the conserved parasite cyt *c*.

### Bacterial reconstitution of cyt *c* hemylation and tests of parasite HCCS specificities

Covalent heme attachment to *c*-type cytochromes requires the IMS protein holocytochrome *c* synthase (HCCS), which binds heme and the N-terminal α-helix of cyt *c* such that the heme vinyl groups are positioned adjacent to the CXXCH of cyt *c* to accelerate stereospecific thioether bond formation (17). Both parasite cyt *c* and *c*-2 retain the N-terminal residues required for HCCS recognition and hemylation (Fig. 3A). Nevertheless, given the unusually high sequence divergence of cyt *c*-2, we considered it uncertain whether this protein would be hemylated by HCCS.

We tested the specificities of *P. falciparum* HCCS and HCC_1_S and evaluated their ability to hemylate parasite cyt *c* and *c*-2. To do so, we cloned and recombinantly co-expressed parasite cyt *c* and *c*-2 with each HCCS homolog in *E. coli* bacteria to reconstitute holocytochrome synthesis in a heterologous in-cell model system. *E. coli* synthesizes heme but lacks cytoplasmic *c*-type cytochromes or holocytochrome *c* biogenesis machinery (27, 28). However, prior studies have shown that heterologous cytoplasmic expression of mitochondrial HCCS and cyt *c* proteins in *E. coli* is sufficient to synthesize holocytochrome *c* (29, 30). Because heme is covalently linked to cyt *c* via stable thioether bonds, the bound heme co-migrates with cyt *c* protein by denaturing SDS-PAGE. After transfer to membrane, cyt *c* hemylation can be detected by exploiting the weak peroxidase activity of bound heme to catalyze luminol chemiluminescence (31). As a positive control, we performed these studies in parallel with cloned, recombinant human HCCS and cyt *c*, which have been characterized in-depth using this in-cell bacterial system (24).

A pGEX vector encoding human HCCS with an N-terminal glutathione-S-transferase (GST) tag (24) was a gift from Robert Kranz (Washington University in St. Louis). We cloned codon-optimized genes for *P. falciparum* HCCS and HCC_1_S into this pGEX vector in frame with the N-terminal GST tag. Human cyt *c* and codon-optimized genes for *P. falciparum* cyt *c* and *c*-2 were cloned into a pET28a vector in frame with an N-terminal hexa-histidine tag. We confirmed expression of all HCCS and cyt *c* proteins in *E. coli* lysates by western blot analysis. Expression of each cyt *c* alone without an HCCS gave no detectable signal by chemiluminescent heme stain, confirming that cyt *c* hemylation depends on HCCS (Fig. 4). Although co-expression of human cyt *c* or malaria parasite cyt *c* or *c*-2 with human (Fig. S3) or parasite HCCS (Fig. 4) resulted in detectable hemylation of all three cyt *c* homologs, co-expression with *P. falciparum* HCC_1_S resulted in no detectable hemylation of human or parasite cyt *c* or *c*-2 (Fig. 4).

**Figure 4.**
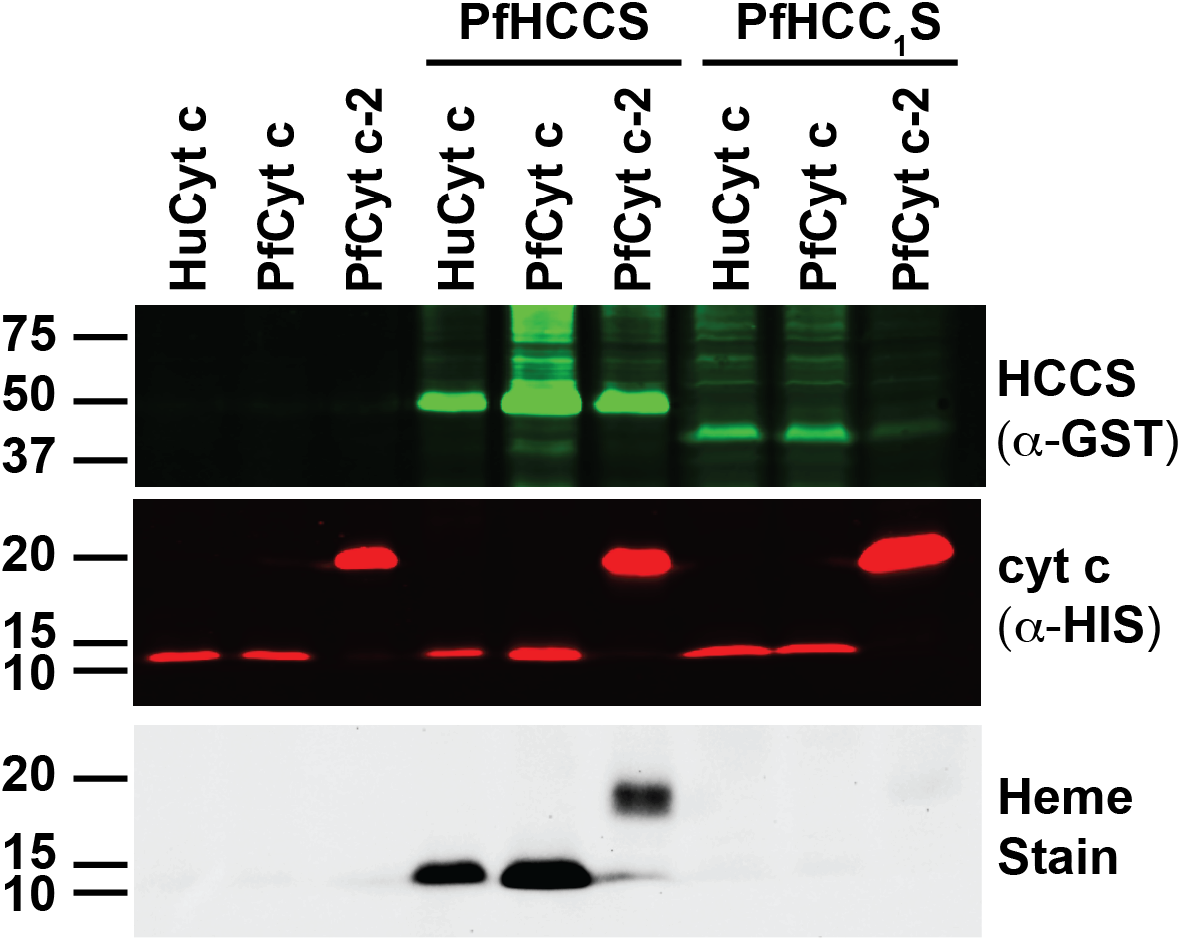
Reconstitution of cyt *c* hemylation in *E. coli*. Each protein was expressed as an N-terminal fusion with GST (HCCS or HCC_1_S) or His_6_ (cyt *c* or *c*-2). Bacterial lysates were fractionated by SDS-PAGE and transferred to nitrocellulose membrane, which was probed with α-GST and α-His-tag antibodies. Bound heme was detected by chemiluminescent heme stain.

We conclude that *P. falciparum* HCCS, but not HCC_1_S, is capable of hemylating parasite cyt *c* and *c*-2, consistent with predicted specificity differences for these HCCS homologs. The ability of *P. falciparum* and human HCCS to promiscuously hemylate cyt *c* homologs from both organisms suggests conserved mechanisms for substrate recognition and heme attachment by these proteins despite their phylogenetic divergence. Co-expression of *P. falciparum* HCC_1_S with parasite cyt *c*_1_ in *E. coli* did not result in detectable cyt *c*_1_ hemylation. However, heterologous reconstitution of mitochondrial cyt *c*_1_ hemylation in *E. coli* has not been reported for any organism, possibly due to cyt *c*_1_ misfolding in *E. coli* and/or a requirement for additional host-cell factors for cyt *c*_1_ hemylation. Nevertheless, the inability of HCC_1_S to hemylate cyt *c* is consistent with a selective role in cyt *c*_1_ biogenesis in parasites, which we are now testing.

### Optical and EPR spectroscopy of cyt *c* homologs reveal differences in axial heme coordination

To assess similarities and differences in the axial heme-coordination states of human cyt *c* and *P. falciparum* cyt *c* and *c*-2, we purified these proteins from *E. coli* lysates using Ni-NTA and size-exclusion chromatography (SEC) and obtained UV-vis absorbance spectra of the oxidized and reduced proteins. All three proteins eluted from the SEC column as a single dominant peak at the expected retention time for a globular monomer (Fig. S4). Spectra of oxidized, ferric (Fe^3+^) human and *P. falciparum* cyt *c* were nearly identical (Fig. 5, A & B) and displayed a Soret peak centered at 410-412 nm and a weaker, broad transition at 535 nm that is characteristic of biaxial His/Met heme coordination in a low-spin complex (32, 33). Reduction by sodium dithionite to the ferrous (Fe^2+^) state red-shifted the Soret peak to 417 nm for both proteins, which also displayed prominent α,β-peaks at ~520 and 550 nm (Fig. 5, A & B), as expected for a hexacoordinate complex (26). The strong spectral similarity between these two proteins is consistent with their high (~60%) sequence identity (Fig. 3A).

**Figure 5.**
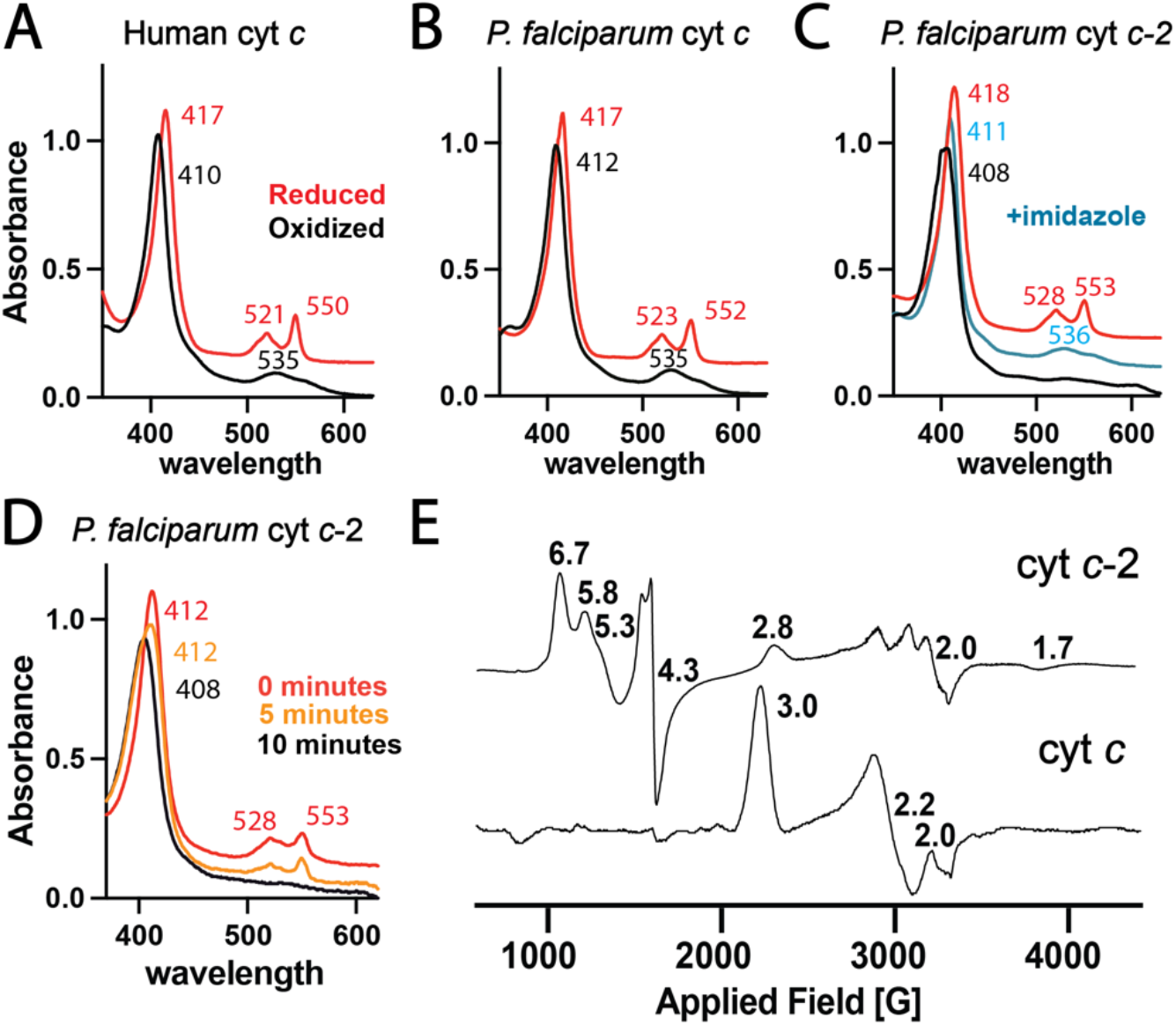
Optical and EPR spectroscopy of cyt *c* homologs. UV-vis absorbance spectra of human cyt *c* (A) and *P. falciparum* cyt *c* (B) and cyt *c*-2 (C). Oxidized and reduced spectra were obtained in 1 mM ammonium persulfate or sodium dithionite, respectively, with 8-12 μM protein. Spectra were normalized to the Soret peak and offset vertically to avoid overlap. For (C), 1 mM imidazole was added to the protein solution. (D) Time-dependent UV-vis absorbance spectra of parasite cyt *c*-2 after addition of sodium dithionite reducing agent. (E) Low-temperature EPR spectra of oxidized *P. falciparum* cyt *c* and *c*-2 acquired at 10 K. EPR g values appear above major features.

In contrast to human and parasite cyt *c*, the spectrum of oxidized, ferric *P. falciparum* cyt *c*-2 displayed a Soret peak at 408 nm, lacked the broad peak centered at 535 nm, and instead featured a weak transition at 602 nm (Fig. 5C). These spectral features suggest that heme is bound in a pentacoordinate, high-spin state with only one axial ligand (32, 33). These unusual properties are consistent with the high sequence divergence in the heme crevice region of cyt *c*-2 compared to human and parasite cyt *c* (Fig. 3A) and the expectation based on prior cyt *c* studies that heme crevice mutations perturb the positioning of Met81 (human numbering) and destabilize its coordination of heme (25, 26).

To further test the provisional conclusion that ferric cyt *c*-2 has only one stable axial ligand, we next probed if this protein could bind exogenous small molecules as a second axial ligand. Incubation of cyt *c*-2 with a stoichiometric excess of imidazole (which mimics the His side-chain) shifted the UV-vis absorbance spectrum to resemble that of ferric human or parasite cyt *c*, including a shifted Soret peak at 411 nm, the loss of a peak at 602 nm, and appearance of a broad transition centered at 536 nm indicative of two axial ligands (Fig. 5C). We observed a similar spectral shift in the presence of hydrogen peroxide (Fig. S5). These spectral changes with exogenous ligands support a structural model that ferric cyt *c*-2 maintains axial His coordination of heme but lacks Met81 ligation (despite retaining Met81) due to structural perturbations from sequence divergence in the heme crevice residues that create an open pocket capable of binding imidazole or hydrogen peroxide as a second axial ligand.

Reduction of cyt *c*-2 with 1 mM sodium dithionite shifted the Soret peak to 418 nm and resulted in α,β-peaks at 528 and 553 nm (Fig. 5C), similar to ferrous human and *P. falciparum* cyt *c* and as expected for biaxial His/Met coordination of heme. The spectrum of reduced cyt *c*-2, however, rapidly reverted over several minutes to that observed for the oxidized ferric state (Fig. 5D). This behavior contrasted with parasite cyt *c*, which remained stably reduced over a similar timescale (Fig. S6). This observation suggests that the heme pocket of cyt *c*-2 is more accessible to oxidation by atmospheric oxygen than parasite cyt *c*, consistent with altered structure and the possibility of enhanced flexibility in the heme crevice region of cyt *c*-2 due to sequence divergence.

To test and define differences in heme coordination structures and spin states of *P. falciparum* cyt *c* and *c*-2, we acquired low-temperature electron paramagnetic resonance (EPR) spectra of the oxidized proteins at 10K. The EPR spectrum of parasite cyt *c* was very similar to previously published spectra for human and yeast cyt *c* (32, 34, 35), with clear signals at g values of 3.0 and 2.2 indicative of a low-spin iron center and as expected for hexacoordinate His/Met ligation (Fig. 5E). This similarity is consistent with the high sequence identity and very similar UV-vis absorbance spectra of these proteins (Fig. 5, A & B).

The EPR spectrum of *P. falciparum* cyt *c*-2, however, differed substantially from human and parasite cyt *c*, with major features at g values of 6.7, 5.8, 5.3, 4.3, 2.8, and 2.0 (Fig. 5E and Fig. S7). This spectrum indicates a predominant population of a high-spin heme center that is consistent with the presence of a single axial ligand, as suggested by the UV-vis absorbance spectrum. These high-spin features are similar to those reported for the H19M mutant of human cyt *c*, which was also proposed to have a single axial ligand (32). The cyt *c*-2 signals at g values of 2.8 and 2.0, however, indicate the presence of a minor low-spin population that may reflect metal contamination in the sample and/or a minor population of either hexacoordinate cyt *c*-2 with biaxial His/Met ligation or a low-lying excited state with pentacoordinate geometry.

### X-ray crystallographic structure of cyt *c*-2 domain-swapped dimer

To better understand the structural features and unusual axial heme environment of cyt *c*-2, we determined the X-ray crystallographic structure of this protein. Sparse-matrix crystallization screens resulted in several crystals at pH 5.5 that diffracted well, and experimental phase determination enabled structure determination and refinement of a 2.3 Å-resolution structure (Table S1). Although cyt *c*-2 is predominantly monomeric in solution at pH 7.5 (Fig. S4), the crystal structures revealed a domain-swapped dimer (DSD) in which the C-terminal α-helix of each monomer had exchanged positions with each other to bind the neighboring subunit (Fig. 6A). This dimeric conformation has been observed for yeast and mammalian cyt *c* under conditions containing ethanol or detergents (36, 37). All conditions that resulted in parasite cyt *c*-2 crystals contained polyethylene glycol (PEG), and crystals formed slowly over several weeks, which may have favored crystallization of the dimeric state. Despite extensive screening, including conditions without PEG or alcohols, we were unable to crystallize a monomeric form of cyt *c*-2. These observations suggest that cyt *c*-2 has local structural heterogeneity, perhaps in the heme crevice region, that disfavors crystallization of the monomer and leads instead to slow crystallization of the DSD.

**Figure 6.**
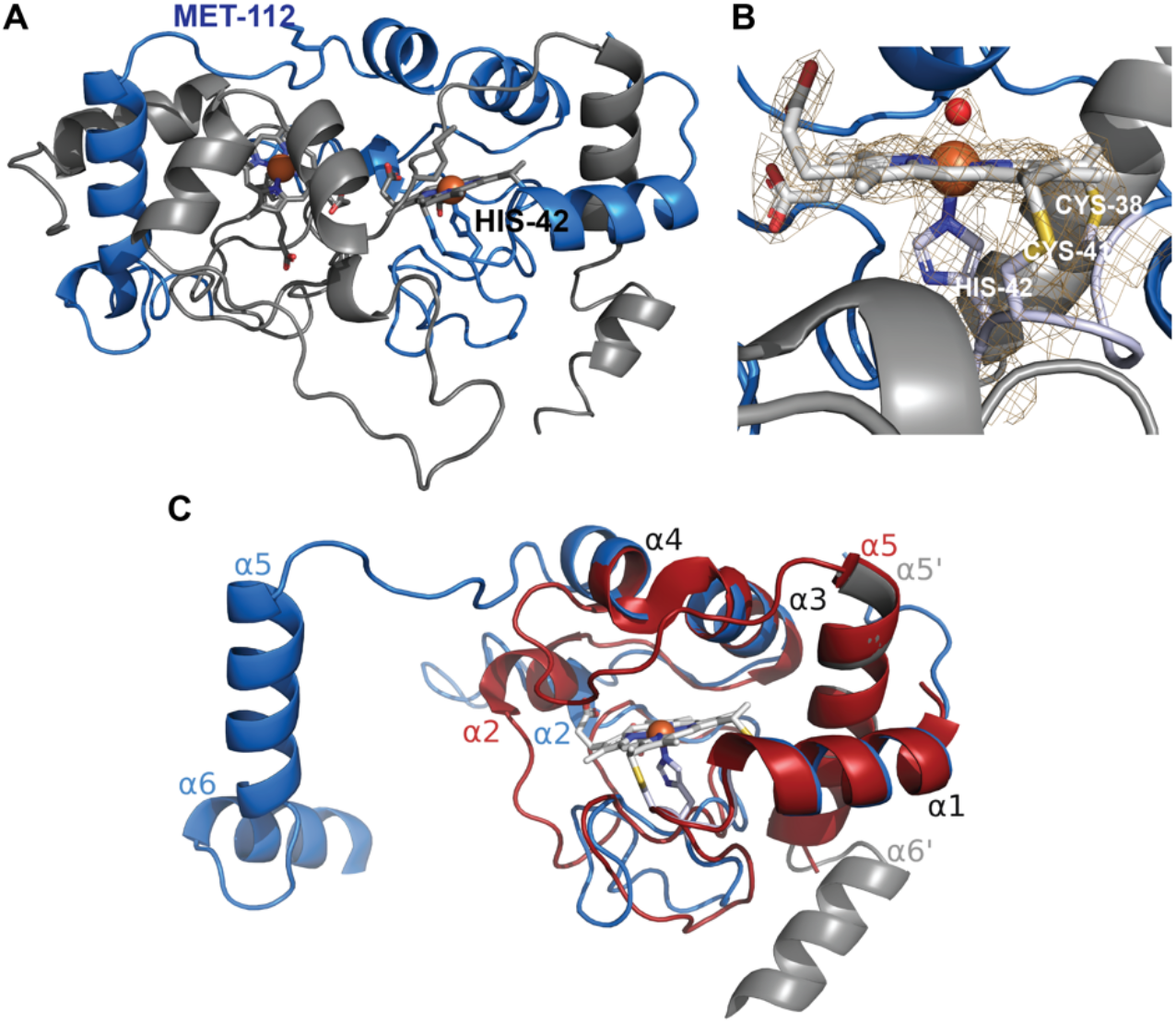
X-ray crystallographic structure of *P. falciparum* cyt *c*-2. (A) 2.3 Å-resolution structure of the domain-swapped dimer (PDB 7TXE), with the two subunits found within the unit cell in blue and gray and His42 and Met112 sidechains shown for one subunit. (B) Active site of one subunit showing bound heme, axial coordination by His42, thioether bonds to Cys41 and Cys38, and 2 |F_o_|-|F_c_| electron-density map (dark orange mesh) contoured at 1.6 σ and showing density for ordered water molecule (red sphere) positioned for axial ligation of heme, in place of Met112. (C) Structural backbone alignment (RMSD = 2.04 Å) of blue subunit from cyt *c*-2 domain-swapped dimer (PDB 7TXE), WT human cyt *c* (red, PDB 3ZCF), and the C-terminal helices (α5’ and α6’) of the second subunit of the cyt *c*-2 dimer (gray). Data collection and refinement statistics are in Table S1.

Despite differences from the solution state, the cyt *c*-2 dimer provided important insights into key structural features of this protein. Despite its unusually high sequence divergence, cyt *c*-2 retains an overall fold typical of eukaryotic cyt *c* proteins (36), including structural similarity in the positions of α-helices 1, 3, and 5 (22). Our structure confirmed the presence of bound heme tethered within the binding pocket by covalent thioether bonds and axial His ligation by the conserved CXXCH motif residues (Fig. 6B). Reciprocal swapping of C-terminal helices mispositioned the conserved Met112 (corresponding to human Met81), disrupting its ability to coordinate the iron of bound heme. Instead, we observed electron density above the iron suggestive of axial coordination by a water molecular, as observed in other cyt *c* DSD structures (36). We also noted several points of structural divergence. First, the 2^nd^ α-helix and residues immediately preceding it are not structurally conserved and adopt a more open conformation that increases the solvent exposure to heme. Second, the N- and C-termini of cyt *c*-2, which contain sequence extensions absent in human cyt *c* (Fig. 3A), adopt random-coil and α-helical conformations, respectively (Fig. 6A). Finally, sequence divergence in cyt *c*-2 that is N-terminal to Met112, including loss of the conserved PGTK sequence found in human and *P. falciparum* cyt *c* and insertion of an unusual His residue (Fig. 3A), results in a longer α-helix 4 relative to human cyt *c* (Fig. 6C). This secondary-structure extension in cyt *c*-2 may disfavor coordination of heme by Met112 and thus increase local structural heterogeneity in this region that enhances solvent access to heme and disfavors crystallization as a monomer.

### Proteomic analysis of parasite cyt *c* and *c*-2 interactions

To identify similarities and differences in protein-interaction partners for parasite cyt *c* and *c*-2 that might clarify their functions, we immunoprecipitated (IP) each endogenously tagged protein and used tryptic digest followed by liquid chromatography-tandem mass spectrometry to identify co-associating proteins. For these experiments, we used CRISPR/Cas9 to introduce a C-terminal HA tag at the endogenous locus for each protein (described below) as an affinity handle for α-HA IP and used untagged parental Dd2 parasites as a negative control. Both cytochromes selectively co-purified with a range of mitochondrial matrix and IMS proteins, including known subunits of the ETC respiratory complexes, and over 60% (23 of 36) of these proteins were identified in both samples (Fig. 7). These results are consistent with the expected IMS localization of both proteins and suggest similar overall protein-protein interactions. Modest differences were observed in the copurifying proteins, but these differences did not strongly differentiate their functions.

**Figure 7.**
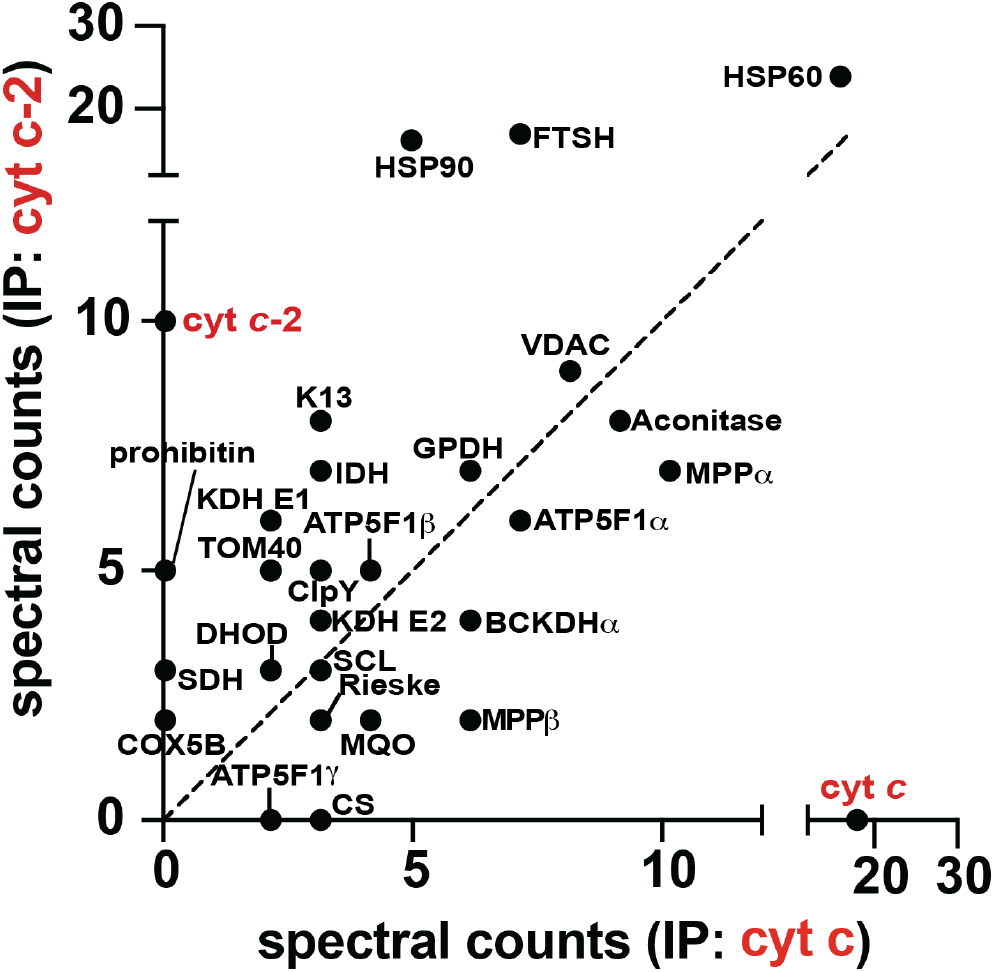
IP/MS studies of protein-protein interactions for blood-stage *P. falciparum* cyt *c* and *c*-2. Bait proteins were immunoprecipitated by anti-HA-tag pull-down, and associated proteins were identified by tryptic digest and tandem mass spectrometry. Spectral counts of detected mitochondrial proteins are shown for each IP sample. None of the detected proteins were identified by anti-HA-tag IP of parental Dd2 parasites, except for HSP60 which was enriched >2-fold relative to Dd2. The dashed line has a slope of one. Source data for mitochondrial and all detected proteins are given in Dataset S1.

Curiously, the kelch-13 protein (PF3D7_1343700) was selectively identified in pull-downs of both cytochromes. This protein is thought to function in the pathway of hemoglobin import into the digestive vacuole, and mutations in its propeller domain underpin parasite resistance to artemisinin (38, 39). Recent work has suggested that kelch-13 can localize to mitochondria and pull down with a variety of mitochondrial proteins, with these interactions appearing to increase upon artemisinin treatment (40). The physical determinants and functional consequences of these associations remain uncertain. Our observation is consistent with this prior work, although the interactions that underpin direct or indirect association of cyt *c* and *c*-2 with kelch-13 remain undefined.

### Critical tests of *c*-type cytochrome importance for blood-stage parasites

The strong sequence conservation and similar physical properties between *P. falciparum* and human cyt *c* suggest that this parasite homolog (PF3D7_1404100) retains a conserved, essential function in the mitochondrial ETC. Indeed, recent genome-wide knockout studies in both rodent and human parasites indicated that this gene is refractory to disruption (41, 42). The divergent sequence and unusual physical properties of *P. falciparum* cyt *c*-2 (PF3D7_1311700), however, seem incompatible with a canonical ETC role and raise substantial uncertainty on its essentiality for blood-stage parasites. Recent genome-wide knockout studies have also given conflicting results for this gene, which was reported to be essential for *P. falciparum* human parasites but fully dispensable for *P. berghei* rodent parasites (41, 42).

To directly test the essentiality of each cyt *c* homolog for blood-stage *P. falciparum* growth and viability, we used CRISPR/Cas9 to tag the endogenous genomic locus for cyt *c*, cyt *c*-2, and cyt *c*_1_ in both Dd2 and NF54 parasites to encode a C-terminal HA-FLAG epitope tag fusion and the aptamer/TetR-DOZI system (43). In this knockdown (KD) system, protein expression is enabled in the presence of anhydrotetracycline (aTc) but tightly repressed after aTc washout. Correct and complete integration into the target genomic locus was confirmed for all three genes by PCR analysis of Dd2 and NF54 parasites and by Southern blot for editing cyt *c*-2 in Dd2 parasites (Fig. S8). Expression of each tagged protein was detected by anti-HA-tag western blot in +aTc conditions, but these proteins were nearly undetectable after three days of growth in -aTc conditions, indicating robust knockdown (Fig. 8, A-C).

**Figure 8.**
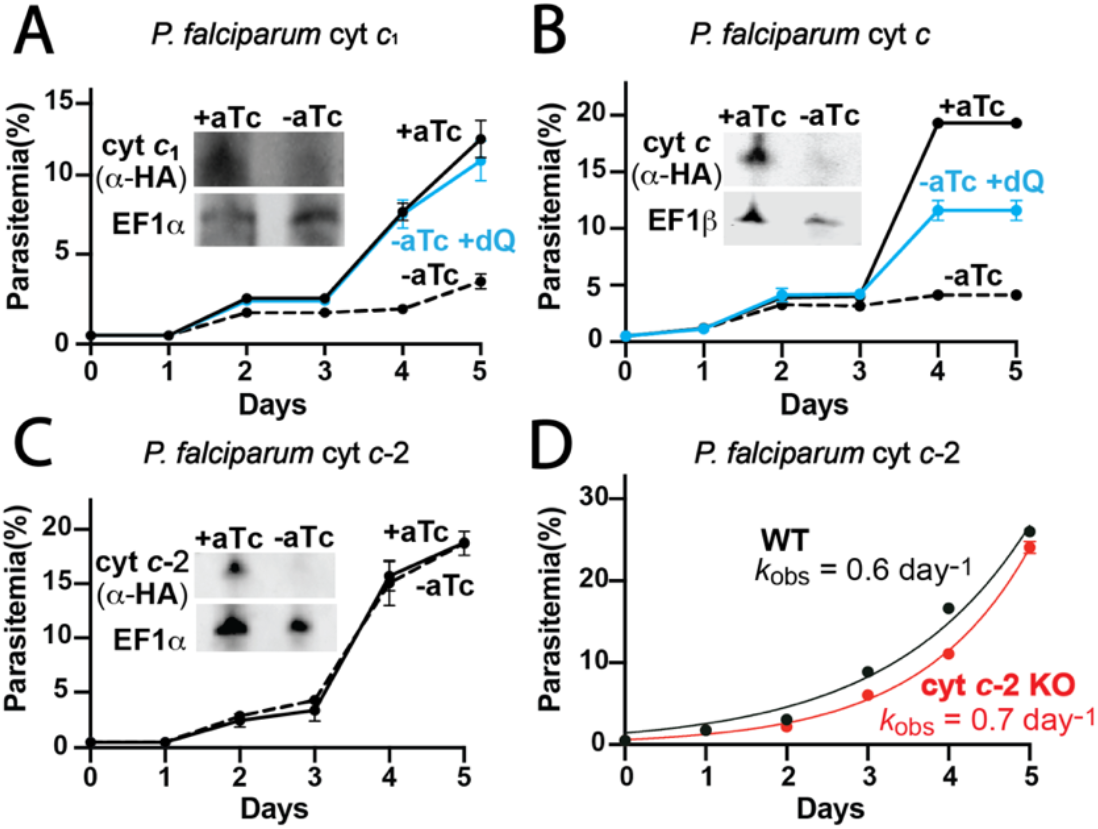
Functional tests of contributions by cyt *c*, *c*-2, and *c*_1_ to blood-stage parasite growth. Synchronous growth assays of Dd2 parasites tagged with the aptamer/TetR-DOZI cassette for inducible knockdown of cyt *c*_1_ (A), cyt *c* (B), and cyt *c*-2 (C), in the presence or absence of 0.5 μM anhydrotetracycline (aTc) and 15 μM decyl-ubiquinone (dQ). Insets are α-HA and α-EF1α or β western blots of parasites lysates harvested on day 3 of the continuous growth assay. (D) Asynchronous growth assay of WT or cyt *c*-2 knockout (KO) NF54 parasites. Data were fit to an exponential growth equation to determine observed rate constants (*k*_obs_). For all growth assays, data points and error bars are the mean and standard deviation of biological triplicates.

All tagged parasites grew normally in synchronous growth assays in the presence of aTc. Washout of aTc, however, resulted in a strong growth defect after 3 days for Dd2 and NF54 parasites containing tagged cyt *c* and *c*_1_ genes (Fig. 8, A and B, and Fig. S9). Blood-smear analysis of these cultures revealed widespread parasite death (Fig. S10). In contrast, aTc washout had no detectable impact on growth of Dd2 or NF54 cyt *c*-2 KD parasites (Fig. 8C and Fig. S9), suggesting that this homolog is not essential for growth of blood-stage *P. falciparum*. To further test this conclusion and to rule out an alternative explanation that KD of cyt *c*-2 is insufficiently stringent to observe a growth phenotype, we stably disrupted the cyt *c*-2 gene in NF54 parasites to create a knockout (KO) line (Fig. S8). The KO parasites grew very similar to parental WT parasites, with a nearly identical rate constant for asynchronous exponential growth (Fig. 8D). This similarity strongly supports the conclusion that the divergent cyt *c*-2 is fully dispensable for blood-stage parasite growth.

The prevailing paradigm posits that oxidative recycling of ubiquinone is the sole essential function of the ETC in blood-stage parasites (10). In direct support of this model, we observed that exogenous addition of oxidized decyl-ubiquinone rescued growth of Dd2 parasites from KD of cyt *c* and *c*_1_ in -aTc conditions (Fig. 8, A and B). To further test and extend this conclusion, we asked if KD of cyt *c* or *c*_1_ sensitized parasites to mitochondrial depolarization by proguanil. This drug targets a secondary pathway in parasites for maintaining mitochondrial transmembrane potential that only becomes essential upon ETC dysfunction (10, 44). Growth of cyt *c* or *c*_1_ KD parasites cultured in +aTc conditions was unaffected by 1 μM proguanil, as expected for an active ETC. However, when 1 μM proguanil was combined with -aTc conditions, decyl-ubiquinone was unable to rescue parasites from lethal growth defect on day four (Fig. 9, A and B). Analysis of parasites on day three of these growth assays by fluorescence microscopy revealed that 1 μM proguanil selectively resulted in dispersed rather than focal staining by MitoTracker Red in cyt *c* or *c*_1_ KD parasites grown in -aTc conditions (Fig. 9, C and D, and Fig. S11), which is indicative of mitochondrial depolarization (10, 15). These observations strongly support our conclusion that loss of cyt *c* or *c*_1_ results in lethal ETC dysfunction that sensitizes parasites to mitochondrial depolarization when treated with proguanil.

**Figure 9.**
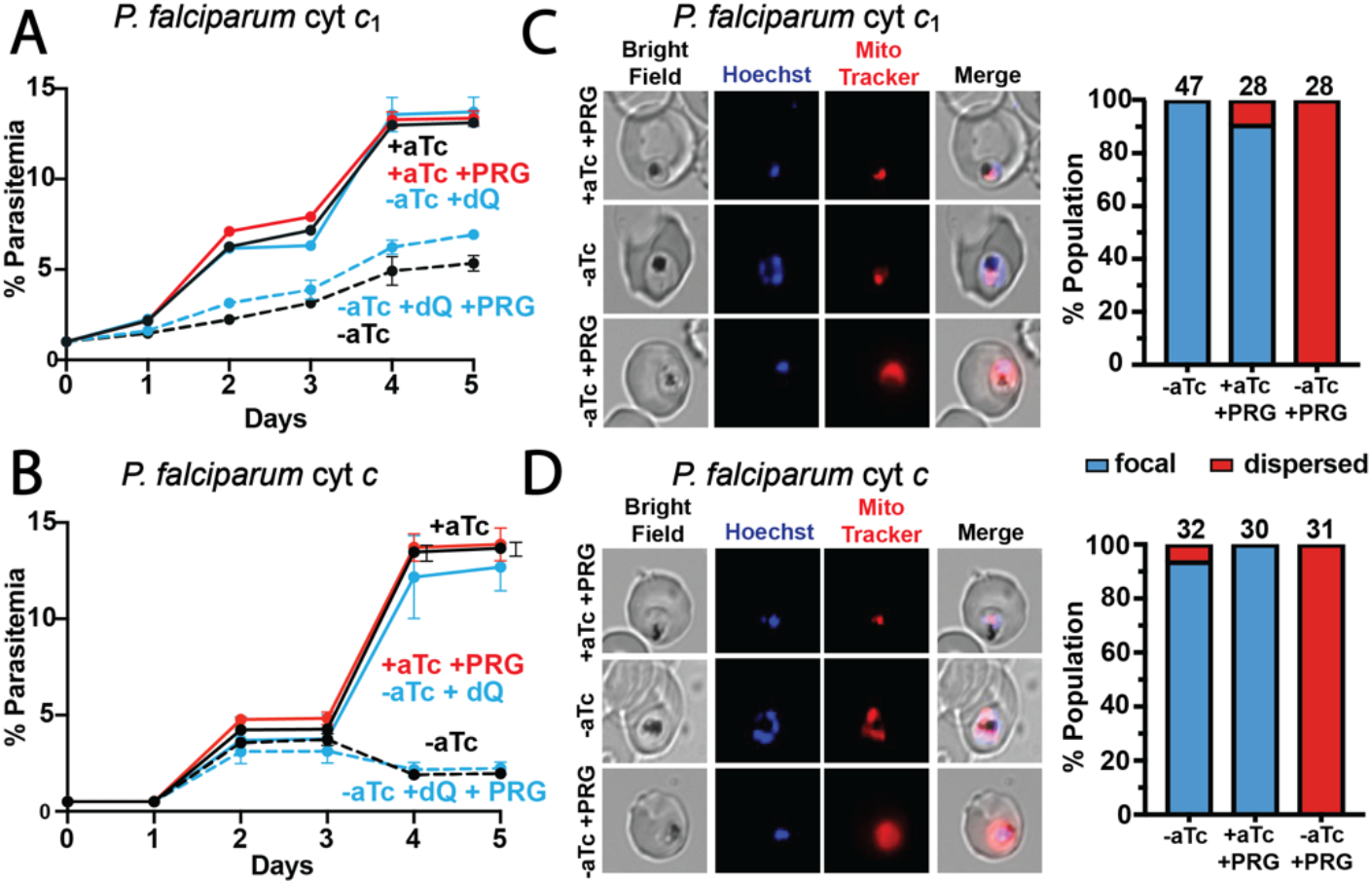
Loss of ETC cytochromes sensitizes parasites to proguanil. Synchronous growth assays of Dd2 parasites tagged with the aptamer/TetR-DOZI cassette for inducible knockdown of cyt *c*_1_ (A) and cyt *c* (B) in the presence or absence of 0.5 μM anhydrotetracycline (aTc), 1 μM proguanil (PRG), and/or 15 μM decyl-ubiquinone (dQ). Data points and error bars are the mean ±SD of biological triplicates. Fluorescence microscopy images of live cyt *c*_1_ (C) or cyt *c* (D) aptamer/TetR-DOZI Dd2 parasites cultured 3 days ±aTc and ± 1 μM proguanil and stained with Hoechst or MitoTracker Red (10 nM). Parasites were imaged in biological duplicate experiments to count 28-47 total parasites, score MitoTracker staining as focal or dispersed, and depict the percentage of the total population under each condition displaying the indicated mitochondrial phenotype.

## DISCUSSION

The mitochondrial ETC of *Plasmodium* parasites has been the subject of intense drug-development efforts over the past several decades to block parasite growth in blood, liver, and mosquito stages (9, 10, 16, 45–48). Nevertheless, key ETC functions have remained uncertain and puzzling. Indeed, many mitochondrial proteins in *Plasmodium* have divergent sequence features and/or are specific to apicomplexan parasites (15, 20, 49–51). In addition, the activity and essentiality of distinct ETC components vary across different parasite developmental stages with the shifting nutritional availability and energy requirements of *Plasmodium* in discrete host environments (1, 11, 13, 20, 48, 52, 53). This functional complexity and the sparsity of direct molecular studies of complexes III and IV have left critical gaps in our basic understanding of parasite ETC function and evolution (3, 54, 55).

### Direct tests of mitochondrial cytochrome functions

We have carried out direct tests of ETC cytochrome functions in *P. falciparum*, focusing on the nuclear-encoded *c*-type cytochromes. Our results indicate that cyt *c*_1_ and the more conserved cyt *c* (PF3D7_1404100) are essential for viability of blood-stage *P. falciparum*, while the divergent cyt *c*-2 (PF3D7_1311700) is dispensable in this stage.

The ability of dQ to rescue parasites from loss of cyt *c* or *c*_1_ provides direct experimental support to the general paradigm, based predominantly on inhibitor studies of cyt *b*, that the essential ETC function in blood-stage parasites is oxidative recycling of ubiquinone to support essential dehydrogenase activity, especially dihydroorotate dehydrogenase (Fig. 1) (9, 10, 14). Prior work has also suggested that parasites have a secondary and redundant pathway for polarizing the inner mitochondrial membrane that is independent of proton translocation by complexes III and IV and can be blocked by low dose proguanil (10, 44). In support of this model, we observed that loss of cyt *c* or *c*_1_ and concomitant defects in ETC-mediated proton translocation did not result in mitochondrial depolarization, which only occurred upon combinatorial treatment with proguanil (Fig. 9). These results are also consistent with our prior study of the mitochondrial acyl carrier protein in *P. falciparum*, whose knockdown results in loss of the Rieske Fe-S protein required for complex III function and can be bypassed by exogenous dQ (15).

Recent proteomic studies of ETC composition in *P. falciparum* and *T. gondii* have identified the mitochondrial processing peptidase (MPP) α and β subunits as key constituents of complex III (20, 56, 57), as observed in plants (58, 59). MPP cleaves the N-terminal leader sequences of proteins imported into mitochondria and appears to be essential for *P. falciparum* (42). We observed that both parasite cyt *c* homologs pulled-down with MPP subunits, with somewhat higher detection of MPP association for cyt *c* than *c*-2 (Fig. 7), consistent with complex III interactions. Whether MPP enzymatic activity requires or is independent of association with complex III is unstudied in parasites. However, our results may provide insight into this question. Loss of cyt *c*_1_ is expected to strongly destabilize complex III, with cyt *c*_1_ knockout resulting in >80% reduction in cyt *b* levels in yeast (60). The ability of dQ to rescue parasites from cyt *c*_1_ knockdown (Fig. 7A and 8A) suggests that MPP activity may persist independent of complex III integrity, since dQ is not expected to bypass more general defects in MPP processing. Biochemical precedent for autonomous MPP function is provided by yeast and humans, in which MPP is a soluble matrix enzyme that functions independent of complex III (61).

### Cytochrome c duplication in *Plasmodium* and comparison to other eukaryotes

Sequencing of the *P. falciparum* genome revealed that parasites encode two copies of cyt *c*, raising questions about the biochemical rationale for this duplication and the functional roles of the distinct paralogs (2, 50, 62). Confusion and uncertainty increased after genome-wide knockout studies gave conflicting assessments of essentiality (41, 42). Although both proteins are expressed (Fig. 7 and Fig. 8), only the conserved cyt *c* is required for *Plasmodium* ETC activity in blood-stage parasites.

Cyt *c*-2 has the most divergent sequence of any known eukaryotic cyt *c*, with functional properties that are not required for ETC activity in blood-stage parasites. These changes in a eukaryotic cyt *c* homolog are fascinating and uncommon, given stringent fitness constraints that disfavor substantial sequence divergence in cyt *c* in order to retain its essential role in mitochondrial respiration (21, 22). Indeed, the strong sequence conservation between cyt *c* from diverse eukaryotes has served as a paradigm for understanding molecular evolution, organismal phylogeny, and the relationships between natural selection and sequence variation (23, 63, 64).

*Saccharomyces cerevisiae* also has two copies of cyt *c*. However, their sequence identity is high (>80%), and their functions appear to be redundant since both copies must be deleted to cause full respiratory deficiency (65, 66). Cyt *c* duplication is also found throughout the phylum Apicomplexa, including *Toxoplasma* and *Babesia*, and in the closely related photosynthetic chromerid, *Vitrella brassicaformis*. These organisms are similar to *Plasmodium*, with one highly conserved homolog of cyt *c* and a second copy with sequence divergence approaching that of *P. falciparum* cyt *c*-2. *Toxoplasma* resembles *Plasmodium* in that the more conserved homolog (TGME49_219750) is essential while the more divergent homolog (TGME49_229420) appears dispensable for tachyzoite growth in fibroblasts (67). Ancestral gene duplication, including whole-genome duplication in *S. cerevisiae* (68), can account for the multiple copies of cyt *c* in these organisms (69). Apicomplexan cyt *c* and *c*-2 may have evolved from specific duplication of an ancestral cyt *c* that occurred at or near the appearance of apicomplexan parasitism >600 million years ago prior to the last common ancestor of *Vitrella, P. falciparum*, and *T. gondii* (70). Different selective pressures presumably resulted either in strong sequence conservation between both copies in yeast or in substantial divergence in Apicomplexa and *Vitrella*.

### Divergent function of *Plasmodium* cytochrome *c-*2

The *P. falciparum* genome is highly reduced, and all genes are expected to enhance parasite fitness at some point in the *Plasmodium* life cycle and/or under specific growth conditions or stresses (62, 71, 72). Retention of the cyt *c*-2 coding sequence in parasites and conservation of its covalent heme-binding features and axial His/Met ligands thus suggests a selective pressure to preserve gene function and to resist sequence degradation, in contrast to cyt *c* pseudogenes found in humans and other organisms that acquired disabling frame-shift and nonsense mutations (69, 73). What then is the molecular function of *Plasmodium* cyt *c*-2, and what fitness contributions explain its retention in the parasite genome?

Our structural and spectroscopic studies of cyt *c*-2 reveal an unusually open active site in which ferric heme is stably coordinated by only a single axial ligand, His42 (Fig. 5 and 6). These structural features create a pocket above the heme into which small-molecule ligands like imidazole can bind. Conservation of Met112 in parasite cyt *c*-2, despite not stably coordinating ferric heme, may reflect a functional role for this residue in transient interactions with heme, including key steps in hemylation by HCCS (24). Destabilization of axial heme coordination by Met112 is expected to perturb the cyt *c*-2 reduction potential. Preliminary measurements indicate a redox potential ~260 mV for parasite cyt *c*, which is very similar to human cyt *c*, but suggest a very different potential for cyt *c*-2 than expected for canonical ETC function (26, 32). Overall, cyt *c*-2 has divergent physical properties that defy a conserved function in the mitochondrial ETC, even if hemylation and mitochondrial localization are retained.

Cyt *c* in mammals and other organisms plays a second critical role in cellular apoptosis, during which cyt *c* binds cardiolipin and transiently adopts an altered conformation that destabilizes axial Met coordination and opens up the heme pocket to enhance intrinsic peroxidase activity (26, 74, 75). Our studies suggest that *P. falciparum* cyt *c*-2 has evolved constitutive physical properties that may mimic aspects of the transient conformational state adopted by canonical cyt *c* homologs during apoptosis, including disruption of heme coordination by Met and an open pocket above heme capable of binding peroxides and other ligands (Fig. S5). Based on these features, we hypothesize that cyt *c*-2 has evolved constitutive peroxidase function that is enhanced relative to cyt *c*, which we are currently testing. This proposed function is notable, since *P. falciparum* lacks canonical heme-dependent peroxidases and catalases typically involved in antioxidant protection (2, 3). Cyt *c*-2 is dispensable for blood-stage parasites but may play a critical role in other host environments or specific growth conditions, including increased oxidative stress. Preliminary experiments, which suggest that cyt *c*-2 KO parasites are 2-3-fold more sensitive to artemisinin than parental NF54 parasites (i.e., lower EC_50_ values), support this hypothesis.

Cyt *c*-2 transcription is substantially upregulated (~80 fold) in mosquito-stage ookinetes relative to blood-stage trophozoites (76, 77), suggesting heightened function and fitness contributions during parasite growth in the insect vector. This observation is intriguing, since oxidative mitochondrial activity and ETC flux in *Plasmodium* increase during parasite development in mosquitoes and may enhance production of hydrogen peroxide and other reactive oxygen species as ETC by-products (11, 53, 78–82). We speculate that cyt *c*-2 may contribute to oxidative metabolism and/or stress protection in this stage, including possible impacts on parasite sensitivity to drugs like atovaquone that target mitochondrial function during growth in mosquitoes (45, 80). We are currently testing cyt *c*-2 function during parasite infection of mosquitoes.

## MATERIALS AND METHODS

### Cloning

For episomal protein expression in *P. falciparum* parasites, the genes encoding cyt *c* (PF3D7_1404100), cyt *c*-2 (PF3D7_1311700), HCCS (PF3D7_1224600), and HCC_1_S (PF3D7_1203600) were PCR-amplified from Dd2 parasite gDNA or cDNA using primer sets 1/2, 3/4, 5/6, and 7/8, respectively (Table S2), and cloned into pTEOE (83) at the Xho1/AvrII sites in frame with a C-terminal GFP tag using ligation-independent methods, as previously described (15). Cloning of cyt c_1_ (PF3D7_1462700) into pTEOE with C-terminal GFP tag was described previously (15). Correct plasmid sequences were confirmed by Sanger sequencing.

For recombinant expression in *E. coli*, codon-optimized genes (most abundant codon in *E. coli* for each amino acid) for human cyt *c* and *P. falciparum* cyt *c*, cyt *c*-2, cyt *c*_1_, HCCS, and HCC_1_S were ordered as gBlocks™ from IDT and cloned by ligation-dependent (*P. falciparum* cyt *c*, primers 9/10) or ligation-independent (cyt *c*_1_, primers 11/12; cyt *c*-2, 13/14; HCCS, 15/16; HCC_1_S, 17/18; Hu cyt *c*, gBlock contained vector-specific sequence) methods into pET28a (cytochromes, NdeI/XhoI sites, in frame with N-terminal His_6_ tag, Novagen 69864) or pGEX (HCCS/HCC_1_S, BamHI/XhoI sites in frame with N-terminal GST tag). Human HCCS in pGEX was a gift from Robert Kranz (24). Plasmid sequences were confirmed by Sanger sequencing.

For disruption of the cyt *c*-2 gene, two different guide RNA sequences corresponding to AGAACAGCAGGAATGAGTAA and AATTCAGGATTTACGAGTTG were cloned by ligation-independent methods using primer pairs 19/20 and 21/22 into a modified pAIO CRISPR/Cas9 vector encoding *Sp*Cas9 and containing a HindIII site for gRNA cloning (15, 84). To disrupt the cyt *c*-2 gene by double-crossover homologous recombination, a repair plasmid was made by ligation-independent cloning (primers 51/52) to insert a gBlock™ fragment ordered from IDT into the XhoI/Avr2 sites of the pPM2GT vector (85) that encodes a hDHFR selection cassette. The gBlock consisted of a 3’ homology flank (containing the last 138 bp of the 3’ end of the cyt *c*-2 coding sequence downstream of the codon for Met112 and 14 bp of the 3’ UTR), an AflII site for linearization, and a 5’ homology flank (containing the last 56 bp of the 5’ UTR plus the first 111 bp of the cyt *c*-2 coding sequence upstream of the sequence encoding the CXXCH motif). Correct plasmid sequences were confirmed by Sanger sequencing.

To tag cyt *c*, cyt *c*-2, cyt *c*_1_ for inducible knockdown using the aptamer/TetR-DOZI system (43), guide RNA sequences corresponding to CTTAATAGAATATTTGAAGA, ACAAAGAAAGGAAAGACAAG, and GCAACTAGAAGAATTGACTT targeting genes for cyt *c*, cyt *c*-2, cyt *c*_1_, respectively, were cloned into the pAIO CRISPR/Cas9 vector using ligation-independent methods (primers 23/24, 27/28, 25/26) by the Mutation Generation & Detection Core at the University of Utah. Donor repair plasmids for each gene were created by using assembly PCR to create a fused sequence containing a 3’UTR homology region (HR) followed by an AflII site and then a 5’ homology region that contained the 3’ end of the coding sequence (without stop codon). This fused insert was then cloned into pMG75 using AscI/AatII cloning site and ligation-independent methods and confirmed by Sanger Sequencing. Donor plasmids to tag each gene were cloned with the following primer sets: cyt *c* (3’ HR: 29/30, 5’ HR: 31/32, including shield mutation), cyt *c*-2 (3’ HR: 37/38, 5’ HR: 39/40), and cyt *c*_1_ (3’ HR: 33/34, 5’ HR: 35/36).

### Parasite culturing

Experiments reported in this study were performed using *P. falciparum* parasite strains Dd2 (86) and NF54 (87). The identities of these strains were confirmed by their expected drug-sensitivity profiles and were *Mycoplasma*-free by PCR test. Parasite culturing was done in Roswell Park Memorial Institute medium (RPM1-1640, Thermo Fisher 23400021) supplemented with 2.5 g/L Albumax I Lipid-Rich BSA (Thermo Fisher 11020039), 15 mg/L hypoxanthine (Sigma H9636), 110 mg/L sodium pyruvate (Sigma P5280), 1.19 g/L HEPES (Sigma H4034), 2.52 g/L sodium bicarbonate (Sigma S5761), 2 g/L glucose (Sigma G7021), and 10 mg/L gentamicin (Invitrogen Life Technologies 15750060). Parasite cultures were maintained at 2% hematocrit in deidentified human erythrocytes obtained from the University of Utah Hospital blood bank, at 37 ̊C, and at 5% O2, 5% CO2, 90% N2. For growth assays with decyl-ubiquinone (dQ, Caymen Chemicals 55486005) and/or proguanil (Sigma 637321), the growth medium was supplemented with 15 μM dQ (DMSO) and/or 1 μM proguanil (DMSO), final concentration. In all cases, the final DMSO concentration was limited to ≤0.3% w/v.

### Parasite transfections

For episomal protein expression using the pTEOE vector, which encodes human dihydrofolate reductase (DHFR) (88) as the drug-selection marker, Dd2 parasites were transfected as previously described (15) by electroporating infected erythrocytes mixed with 1x cytomix containing 50-100 μg midi-prep DNA of purified plasmids and 25 μg of the pHTH transposase plasmid (89) in 0.2 cm cuvettes using a Bio-Rad Gene Pulser Xcell system (0.31 kV, 925 μF). Transfected parasites were allowed to recover in drug-free media for 48 hours and then selected with 5 nM WR 99210 (Jacobus Pharmaceuticals). Transfections involving the pTYEOE vector, which contains yeast dihydroorotate dehydrogenase (DHOD) as the selectable marker (43), were performed with selection in 2 μM DSM1.

For CRISPR/Cas9-based genome-editing experiments to tag cyt *c* (PF3D7_1404100), cyt *c*_1_ (PF3D7_1462700), or cyt *c*-2 (PF3D7_1311700) with a C-terminal HA-FLAG epitope tag and the 10X aptamer/TetR-DOZI system, Dd2 or NF54 parasites were transfected with 50-100 μg each of the linearized pMG75 donor plasmid (with gene-specific homology arms) and pAIO plasmid (84) (encoding Cas9 and gene-specific gRNA) and selected with 6 μM blasticidin-S in the presence of 0.5 μM anhydrotetracycline (Caymen Chemical 10009542). Polyclonal parasites that returned from transfection were genotyped by PCR using primer sets 43/46 and 43/42 (cyt *c*), 44/47 and 44/42 (cyt *c*_1_), and 45/48 and 45/42 (cyt *c*-2) to test for remnant unmodified locus and successful integration, respectively. For cyt *c*-2, parasites were also genotyped by Southern blot, as previously described (90), with digestion of gDNA by Nde1 and hybridization of an oligonucleotide probe created by PCR using primers 41/39. Since PCR and Southern blot analyses indicated full genomic integration without evidence for unmodified genetic loci, polyclonal parasites were used for all assays.

To stably disrupt the cyt *c*-2 gene, NF54 parasites were transfected with 50 μg each of the cyt *c*-2 KO donor plasmid and pAIO CRISPR/Cas9 plasmid (with cyt *c*-2-specific gRNA) and selected with 5 nM WR99210. Polyclonal parasites that returned from transfection were genotyped by PCR using primer sets 45/48 and 49/50 to test for the presence of the WT and modified locus, respectively. This analysis and the qPCR analysis below revealed complete genomic integration without evidence of unmodified cyt *c*-2 gene in the polyclonal culture.

### qPCR analysis of cyt *c*-2 KO parasites

Genomic DNA from NF54 WT, polyclonal cyt *c*-2 KO transfection, or clonal cyt *c*-2 KO parasites was isolated with the Qiagen Blood Miniprep kit and quantified on a NanoDrop™ One^c^ spectrophotometer. To create calibration curves for the WT or disrupted cyt *c*-2 locus, 0.1 – 100 ng total DNA for the WT NF54 or clonal cyt *c*-2 KO sample was added to PowerUP SYBR Green Master Mix (Invitrogen) and amplified in duplicate using primer sets specific to the 5’ and 3’ ends of the WT (5’, primers 53/55; 3’, primers 57/58) or disrupted locus (5’, primers 53/54; 3’, primers 56/58) on a Quantstudio3 Real Time PCR system. The observed C_t_ value for each reaction was plotted as a function of input DNA amount (ng) and fit with a linear function. Genomic DNA from polyclonal cyt *c*-2 KO parasites was assessed via qPCR following the same procedure (four input DNA concentrations, both 5’ and 3’ amplicons, and duplicate samples for 16 total reactions that each targeted the WT or disrupted locus). The abundance of WT or disrupted locus in the polyclonal cyt *c*-2 KO was quantified by applying the standard curve formulas above the observed C_t_ value. For each input DNA amount, the % abundance of the disrupted locus was determined according to the formula: % abundance^KO^ = 100*(ng^KO^)/(ng^KO^ + ng^WT^). The 16 individual % abundance values determined in this fashion for the WT or disrupted locus in the polyclonal cyt *c*-2 or parental NF54 samples were plotted, along with the average ± SD.

### Parasite growth assays

For synchronous growth assays involving aptamer/TetR-DOZI-tagged parasites, asynchronous cultures were treated with 5% D-sorbitol (Sigma S7900) to produce ring-stage parasites. Sorbitol-treated cultures were washed three times with RPMI media lacking anhydrotetracycline (aTc) and then divided into two equal parts before supplementing one part with 0.5 μM aTc. Parasites were diluted to 0.5% parasitemia ±aTc, and cultures were allowed to expand over several days with daily media changes. For asynchronous growth assays of WT and cyt *c*-2 KO NF54 parasites, cultures were diluted to 0.5-1% parasitemia and allowed to expand over several days. For all growth assays, parasitemia was monitored daily by flow cytometry by diluting 10 μL of each parasite culture well from each of three biological replicates into 200 μL of 1.0 μg/mL acridine orange (Invitrogen Life Technologies A3568) in phosphate buffered saline (PBS) and analysis on a BD FACSCelesta system monitoring SSC-A, FSC-A, PE-A, FITC-A, and PerCP-Cy5-5A channels. Daily parasitemia measurements for asynchronous cultures were plotted as a function of time and fit to an exponential growth equation using GraphPad Prism 9.0.

### Immunoprecipitation experiments

Dd2 parasites expressing endogenously tagged cyt *c* or *c*-2 with C-terminal HA-FLAG tags were harvested from ~50 mL of culture by centrifugation, treated with 0.05% saponin (Sigma 84510) in PBS for 5 min at room temperature to lyse erythrocytes, and spun down at 4000 rpm for 30 min at 4°C. Saponin pellets were lysed by resuspension in 1% Triton X-100 in PBS plus protease inhibitor (Thermo Scientific A32955), dispersed by brief sonication on a Branson sonicator equipped with a microtip, incubated for 1 hour on a rotator, and spun at 13,000 rpm for 10 min. The clarified lysates were mixed with equilibrated resin from 30 μl of Pierce anti-HA-tag magnetic beads (ThermoScientific 88836) and incubated for 1 hour at 4°C on a rotator. Beads were placed on a magnetic stand, supernatants were removed by aspiration, and beads were washed 3x with cold TBST (1x TBS supplemented with 0.5% Tween-20). Bound proteins were eluted with 100 μl of 8M urea (in 100 mM Tris-HCl at pH 8.8). Proteins were precipitated by adding 100% trichloroacetic acid (Sigma 76039) to a final concentration of 20% v/v and incubated on ice for 1 hour. Tubes were spun at 13,000 rpm for 25 min at 4°C, supernatants were removed by aspiration, and the protein pellet was washed once with 500 μl of cold acetone. The protein pellets were air-dried for 30 min and stored at −20°C. These samples were then analyzed by mass spectrometry as indicated below.

### SDS-PAGE and Western blot analyses

For analysis of cyt *c* hemylation in *E. coli*, BL21/DE3 bacteria expressing human or *P. falciparum* HCCS or HCC_1_S (encoded by pGEX vector with N-terminal GST tag) and human or parasite cyt *c* or *c*-2 (encoded by pET28a vector with N-terminal His_6_ tag) were grown at 37°C in LB media until reaching an optical density at 600 nm of 0.5, induced with 1 mM IPTG (GoldBio I2481C50) and 100 μM 5-aminolevulinic acid, and grown overnight at 20°C. Bacteria were harvested by centrifugation, washed, and resuspended with 1X TBS (50 mM Tris-HCl, pH 7.4, 150 mM NaCl) supplemented with 1% Triton X-100 v/v and protease inhibitor cocktail (ThermoFisher J61852.XF). Bacteria were lysed by sonication (20 pulses, 50% duty cycle, 50% power) on a Branson sonicator equipped with a microtip and then incubating at 4°C on a rocker for 1 hour. Lysates were clarified by centrifugation (10 min at 14,000 rpm), and total protein was quantified by the Lowry method. 50 μg of total protein was dissolved in 1x SDS sample buffer, heated at 95°C for 10 minutes, fractionated by SDS-PAGE using 12.5% acrylamide gels run at 120V in the Bio-Rad mini-PROTEAN electrophoresis system, and transferred to nitrocellulose membrane for 1 hour at 100 V using the Bio-Rad wet-transfer system. For evaluation of cyt *c* hemylation, membranes were developed for 5 min with the Prometheus™ ProSignal Femto ECL reagent (Genesee Scientific 20-302) and imaged with 10 sec exposure using a BioRad ChemiDoc MP Imaging System.

For evaluation of cyt *c* and HCCS protein expression via western blot, membranes were blocked with 5% (w/v) skim milk in 1x TBS for 1 hour at ambient temperature. Membranes were probed at 1:1000 dilution with mouse monoclonal anti-GST-DyLight800 conjugated antibody (ThermoFisher MA4-004-D800) and mouse monoclonal anti-His_6_ tag-DyLight680 conjugated antibody (ThermoFisher MA1-21315-D680), washed in 1x TBS supplemented with 0.5% (v/v) Tween-20, and imaged on a Licor Odyssey CLx.

To evaluate expression of endogenous cyt *c*, cyt *c*_1_, and cyt *c*-2 in blood-stage *P. falciparum* parasites tagged with the aptamer/TetR-DOZI system and C-terminal HA-FLAG tag, parasites were synchronized with 5% D-sorbitol to rings, grown for 72 hours ±aTc, and harvested by saponin lysis as second-cycle trophozoites and schizonts. Saponin pellets were solubilized in reducing 1X SDS sample buffer, heated for 10 min at 95°C, and clarified by centrifugation for 10 min at 13,000 rpm. Samples were then fractionated by 10% SDS-PAGE and transferred to nitrocellulose membrane as above. Membranes were blocked at room temperature for 1 hour in 1% casein/PBS and probed overnight at 4°C with a 1:1000 dilution of Roche monoclonal 3F10 rat anti-HA-tag antibody (Sigma 11867423001) and 1:1000 dilution of rabbit anti-*P. falciparum* EF1α or EF1β polyclonal antibodies (91). Membranes were washed 3 times, probed with 1:10,000 dilutions of donkey anti-rat DyLight800 2° antibody (Life Technologies SA5-10032) and donkey anti-rabbit DyLight680 2° antibody (Life Technologies SA5-10042), and imaged on a Licor Odyssey CLx.

### Fluorescence microscopy

For live-cell experiments with GFP-tagged cytochromes or HCCS proteins, parasite nuclei were visualized by incubating samples with 1-2 μg/mL Hoechst 33342 (Thermo Scientific Pierce 62249) for 10-20 minutes at room temperature. Mitochondria were visualized by incubating parasites with 10 nM MitoTracker Red CMXROS (Invitrogen Life Technologies M7512) for 15 minutes prior to wash-out and imaging. For mitochondrial depolarization experiments with cyt *c* and *c*_1_ knockdown parasites, cultures were synchronized with 5% D-sorbitol, cultured ±aTc and ±1 μM proguanil for 72 hours before addition of MitoTracker, and imaged as second-cycle trophozoites/schizonts. 28-47 total parasites for each condition across two independent experiments were scored for focal or dispersed MitoTracker signal. For immunofluorescence assay (IFA) experiments, parasites were fixed, stained, and mounted, as previously described (83, 92). For IFA studies, the parasite mitochondrion was visualized using a polyclonal Rabbit anti-HSP60 antibody (Novus NBP2-12734) and AlexaFluor 647-conjugated Goat anti-Rabbit 2° antibody (Invitrogen Life Technologies A21244), the nucleus was stained with ProLong Gold Antifade Mountant with DAPI (Invitrogen Life Technologies P36931), and the endogenously C-terminal HA-tagged cytochromes were visualized with a Roche Rat anti-HA monoclonal 3F10 primary antibody and FITC-conjugated Donkey anti-Rat 2° antibody (Invitrogen Life Technologies A18746). Images were taken on DIC/brightfield, DAPI, GFP, and RFP channels using an EVOS M5000 imaging system, a Nikon widefield fluorescence microscope, or an Airyscan LSM800 confocal microscope (Carl Zeiss). Fiji/ImageJ was used to process and analyze images. All image adjustments, including contrast and brightness, were made on a linear scale. Two-dimensional intensity plots were generated in Fiji/ImageJ with the “plot profile” function using a shared region of interest (identified in images by a white line) on each channel. Maximum intensity projections for Airyscan images were generated in ImageJ using the “z-stack maximum intensity” function.

### Recombinant protein expression and purification

For heterologous expression of hemylated cyt *c*, plasmids encoding *P. falciparum* cyt *c* or cyt *c*-2 with an N-terminal hexahistidine tag (pET28a) and human HCCS with N-terminal glutathione S-transferase (GST) tag (pGEX) were co-transformed into BL21 DE3 *E. coli* cells and selected using 30 μg/ml kanamycin and 50 μg/ml carbenicillin. A saturated overnight culture of transformed bacteria was used to seed 1 L volumes of ZY autoinduction media that were grown at 37° C for 24 hours. Cells were harvested by centrifugation and resuspended in lysis buffer (500 mM NaCl, 20 mM imidazole, 20 mM sodium phosphate, pH=7.5) and a cocktail of protease inhibitors (pepstatin A, leupeptin hemisulfate, aprotinin from Goldbio; phenylmethylsulfonyl fluoride from Acros Organics), bovine pancreas DNAase I (Goldbio), and egg-white lysozyme (Goldbio). After a 20 min incubation at 4° C, the resuspended cells were lysed by sonication (Branson) using 3 x 2 min rounds of sonication with a 33% duty cycle at 75% power and 5 min. incubation on ice between rounds. The lysed sample was then clarified by centrifugation for 1 hour at 18,000 RPM (41,600 x g). The supernatant was collected and incubated with 4 ml of Ni-NTA agarose (Qiagen) for 2 hours at 4° C. The resulting Ni-NTA agarose was loaded into a gravity column and washed with 100 mL of lysis buffer. Bound protein was eluted in 30 ml using 500 mM NaCl, 500 mM imidazole, 20mM sodium phosphate, pH=7.5. The resulting eluate was dialyzed using 12-14 kDa MWCO membrane to remove imidazole using 1x phosphate buffered saline (pH 7.4). After 2 hours, 25 units of thrombin (Sigma) were added to remove the N-terminal His_6_-tag, and the sample was dialyzed a further 12-18 hours at 20°C in 1x PBS. To remove any uncleaved protein that retained the His-tag, the digested protein was separated by a second incubation with Ni-NTA agarose. The unbound flow-through (containing protein lacking the His-tag) was then dialyzed into 100 mM NaCl, 20 mM potassium phosphate, pH=6, prior to loading onto a 5-ml HiTrap SP cation-exchange column (Cytiva) using an Akta Go FPLC system. The SP column was eluted using a linear gradient of 100% buffer A (100 mM NaCl, 20 mM sodium phosphate, pH=6) to 100% buffer B (1 M NaCl, 20 mM sodium phosphate, pH=6) over 100 mL, while collecting 5-ml fractions and monitoring UV absorbance at 280 nm. The protein-containing fractions with cyt *c* or *c*-2 were pooled and injected over an S-100 size-exclusion column (Cytiva) that was eluted with an isocratic flow of 1x PBS at 1 ml/min. Protein-containing fractions were pooled and concentrated to 10 mg/ml using 10 kDa MWCO spin concentrators (Sartorius). Protein purity and identity were confirmed by detection of a single band by Coomassie-stained SDS-PAGE and by LC-MS/MS to confirm the expected sequence.

### Protein crystallization and X-ray structure determination

Crystals of *Plasmodium falciparum* cyt *c*-2 containing bound heme were grown by sitting drop vapor-diffusion method at 21° C using the main-peak fractions from size-exclusion chromatography corresponding to monomeric protein. Crystallization conditions spanned three ratios (1:2, 1:1, and 2:1) in the mixture of protein (10 mg/mL, 10 mM Tris, 80 mM NaCl, 5% glycerol, pH 8) and two different crystallization solutions. The crystal used to determine the initial 2.6 Å-resolution structure was grown in 200 mM sodium chloride, 100 mM Bis-Tris, pH 5.5, 25% PEG 3350. The crystal that diffracted to 2.3 Å resolution was grown in 200 mM ammonium acetate, 100 mM Bis-Tris, pH 5.5, 25% PEG 3350. Crystals were transferred to the crystallization solution (supplemented with 20% glycerol as a cryo-protectant) and subsequently frozen in liquid N_2_. Crystals were maintained at 100K during data collection with the use of a cold nitrogen gas stream.

Diffraction data for the initial 2.6 Å-resolution structure were collected at beamline 12-1 at the Stanford Synchrotron Radiation Lightsource (SSRL). Initial phases for this structure of the *P. falciparum* cyt *c*-2 domain-swapped dimer were determined by single-wavelength anomalous dispersion (SAD) phasing methods in PHENIX using the anomalous signal from the iron in the bound heme. Diffraction data for the 2.3 Å-resolution structure of the domain-swapped dimer of *P. falciparum* cyt *c*-2 were collected at 100K at beamline 9-2 of SSRL and phased by molecular replacement with PHASER in PHENIX using the coordinates from the 2.6 Å-resolution structure as a search model. COOT (93) and PHENIX (94) were used for model rebuilding and refinement, respectively. Protein structures were visualized and aligned in PyMOL, version 2.0 (Schrödinger, LLC). Structure factors and model coordinates were deposited in the RCSB Protein Data Bank under accession codes 7TXE (2.3 Å-resolution) and 7U2V (2.6 Å-resolution). The structural alignment in Fig. 6C was made by aligning residues 25-38 and 71-108 of subunit 1 of parasite cyt *c*-2 (PDB 7TXE) with residues 1-14 and 50-75 of human cyt *c* (PDB 3ZCF) and aligning residues 121-136 of subunit 2 of parasite cyt *c*-2 with residues 87-104 of human cyt *c*.

### UV/Vis absorbance spectroscopy

UV-visible absorbance spectra of ~10 μM protein samples were acquired in phosphate buffered saline (pH 7.4) at 20° C on a Thermo Scientific™ NanoDrop™ One^c^ spectrophotometer with 1-cm cuvette. Protein samples were treated with 1 mM sodium dithionite or ammonium persulfate to obtain reduced or oxidized spectra, respectively.

### EPR Spectroscopy

Electron paramagnetic resonance (EPR) spectroscopy experiments were performed at the Utah State University Department of Chemistry and Biochemistry. For perpendicular mode continuous wave (CW) X-Band EPR measurements, ~100 μL of 100 μM samples were flash frozen in 3 mm inner diameter quartz EPR tubes in liquid nitrogen. CW EPR measurements were performed on a Bruker EMX Plus EPR spectrometer (Bruker Biospin, Billerica, MA) equipped with a liquid helium cryostat operating at X-band (9.38 GHz microwave frequency) with 4000 G sweep width, 10.0 mG modulation amplitude, and 1 mW incident microwave power. Spectra were collected at 5.6-5.7 K and 9.94 K and averaged over 5 scans.

### Mass spectrometry of parasite immunoprecipitation samples

Protein samples isolated by anti-HA-tag IP of endogenous cyt *c* or *c*-2 were reduced and alkylated using 5 mM Tris (2-carboxyethyl) phosphine and 10 mM iodoacetamide, respectively, and then enzymatically digested by sequential addition of trypsin and lys-C proteases, as previously described (95). The digested peptides were desalted using Pierce C18 tips (Thermo Fisher Scientific), dried, and resuspended in 5% formic acid. Approximately 1 μg of digested peptides was loaded onto a 25-cm-long, 75-μm inner diameter fused silica capillary packed in-house with bulk ReproSil-Pur 120 C18-AQ particles, as described previously (96). The 140-min water-acetonitrile gradient was delivered using a Dionex Ultimate 3,000 ultra high-performance liquid chromatography system (Thermo Fisher Scientific) at a flow rate of 200 nl/min (Buffer A: water with 3% DMSO and 0.1% formic acid, and Buffer B: acetonitrile with 3% DMSO and 0.1% formic acid). Eluted peptides were ionized by the application of distal 2.2 kV and introduced into the Orbitrap Fusion Lumos mass spectrometer (Thermo Fisher Scientific) and analyzed by tandem mass spectrometry. Data were acquired using a Data-Dependent Acquisition method consisting of a full MS1 scan (resolution = 120,000) followed by sequential MS2 scans (resolution = 15,000) for the remainder of the 3-s cycle time. Data was analyzed using the Integrated Proteomics Pipeline 2 (Integrated Proteomics Applications, San Diego, CA). Data were searched against the protein database from *P. falciparum* 3D7 downloaded from UniprotKB (10,826 entries) on October 2013. Tandem mass spectrometry spectra were searched using the ProLuCID algorithm followed by filtering of peptide-to-spectrum matches by DTASelect using a decoy database-estimated false discovery rate of <1%.

### Mass spectrometry of purified recombinant protein samples

After purification of recombinant *P. falciparum* cytochrome *c* and *c*-2 from *E. coli* (described above), protein identities were confirmed by proteolytic digestion and tandem mass spectrometry using the Mass Spectrometry and Proteomics Core at the University of Utah, as previously described (15).

## ACKNOWLEDGEMENTS

We thank Seyi Falekun for help with IP/MS experiments; Megan Okada for graphical figure preparation; Robert Kranz for plasmids encoding human HCCS and cyt *c* and helpful discussions; Shelly Minteer for assistance with redox potential measurements; and Bruce Bowler, Amit Reddi, Jared Rutter, Akhil Vaidya, and Dennis Winge for helpful discussions. Research reported in this publication was supported by a Burroughs Wellcome Fund Career Award at the Scientific Interface (to PAS); NIH grants R35GM133764 (to PAS) and R01GM089778 (to JAW); a pilot award (to PAS) from the Utah Center for Iron and Heme Disorders (U54DK110858); and a seed grant (to PAS) from the University of Utah Vice President for Research. PAS is a Pew Biomedical Scholar, supported by The Pew Charitable Trusts. TJSE and AM were supported by NIH training grant T32DK007115. SN was supported by the Utah Native American Research Internship (R25HL108828) and by NIH diversity supplement R35GM133764-04S1. DNA synthesis and sequencing, fluorescence microscopy, creation of CRISPR/Cas9 reagents, recombinant protein mass spectrometry, and flow cytometry were performed using University of Utah core facilities. EPR experiments were performed at Utah State University, with instrumentation supported by NSF grant MRI-0722849. Use of the Stanford Synchrotron Radiation Lightsource is supported by the U.S. Department of Energy Contract No. DE-AC02-76SF00515 and by the NIH (P30GM133894).

## SUPPORTING INFORMATION

**Figure S1.**
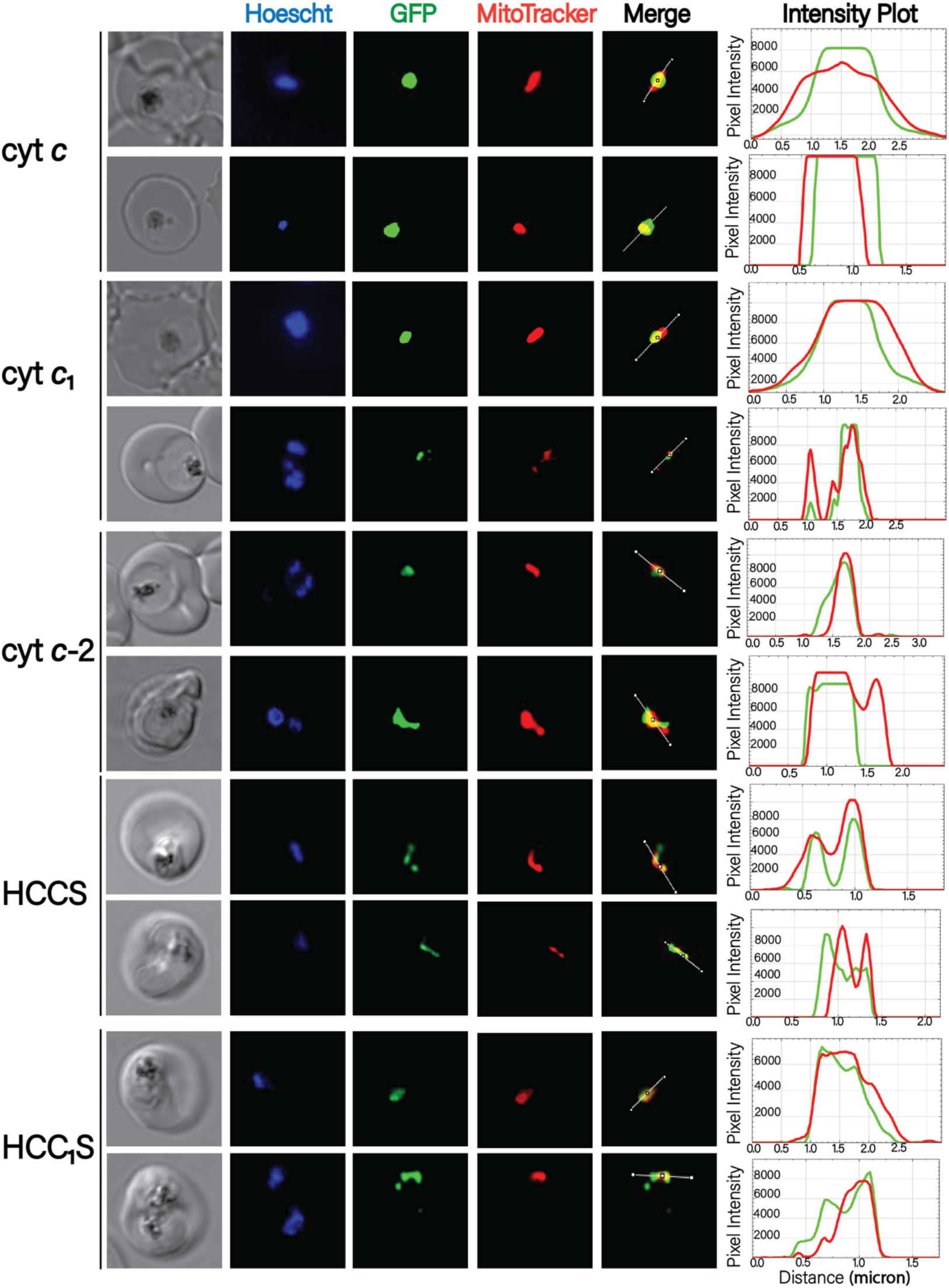
Additional fluorescence microscopy images localizing *c*-type cytochromes and holocytochrome c synthase proteins in blood-stage parasites. Images were obtained by live epifluorescence microscopy of parasites expressing episomal second copies of each protein with a C-terminal GFP tag and stained with MitoTracker Red (mitochondrion) and Hoescht (nucleus). The intensity plot to the right of each image series displays the overlap in pixel intensity for GFP and MitoTracker Red as a function of distance along the white line in the merged image.

**Figure S2:**
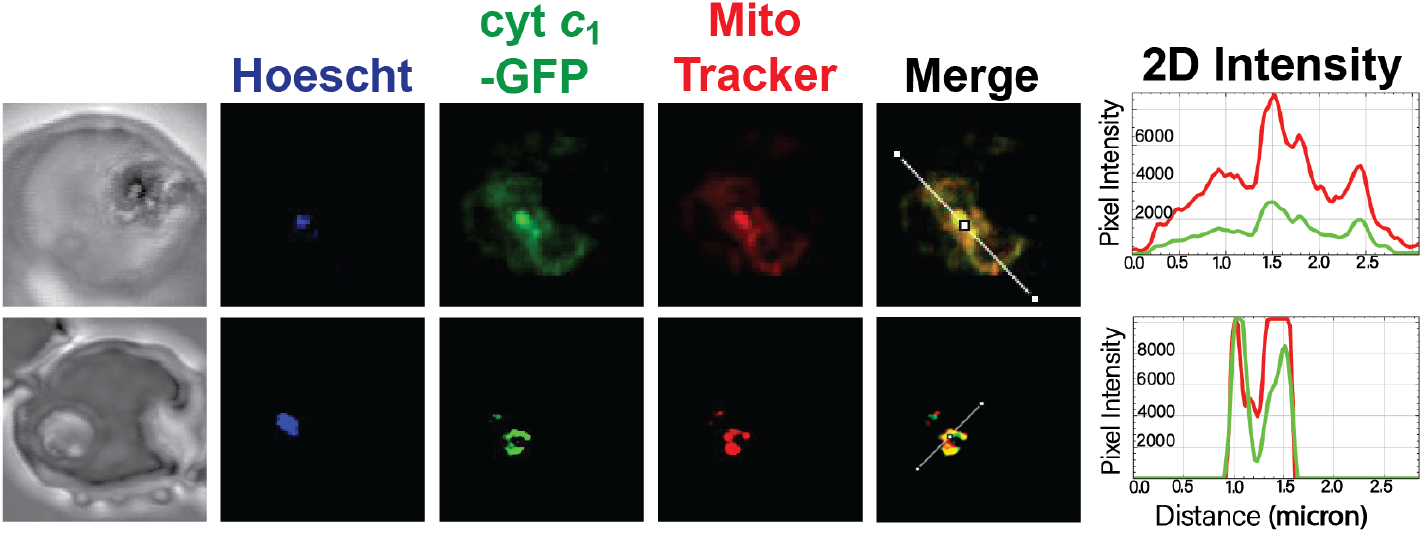
Airyscan confocal fluorescence microscopy of live Dd2 parasites expressing cyt *c*_1_-GFP and stained with 10 nM MitoTracker Red (mitochondrion) and Hoescht (nucleus). Fluorescence images are 2D maximum-intensity projections of z-stack images. The intensity plot to the right of each image series displays the overlap in pixel intensity for GFP and MitoTracker Red as a function of distance along the white line in the merged image.

**Figure S3.**
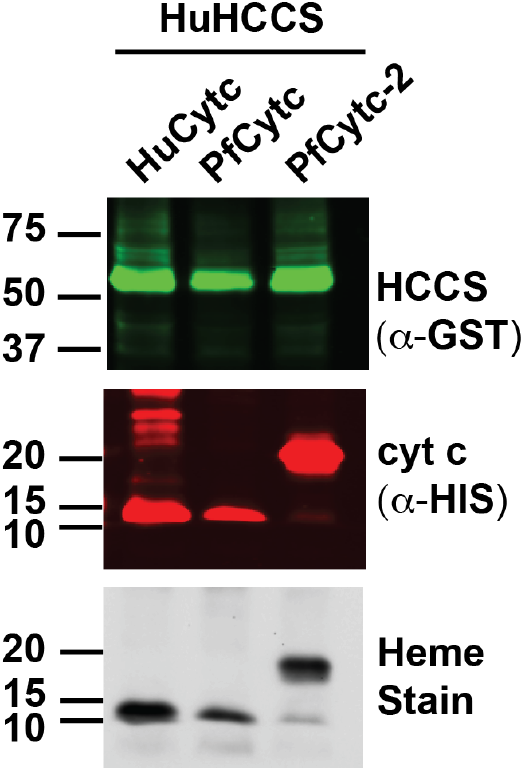
Hemylation of human and *P. falciparum* cyt *c* by human HCCS. Cytochromes and HCCS were expressed as N-terminal fusions with His_6_ and GST, respectively. Bacterial lysates were fractionated by SDS-PAGE and transferred to nitrocellulose membrane, which was probed with α-GST and α-His-tag antibodies. Covalently bound heme was detected by chemiluminescent heme stain.

**Figure S4.**
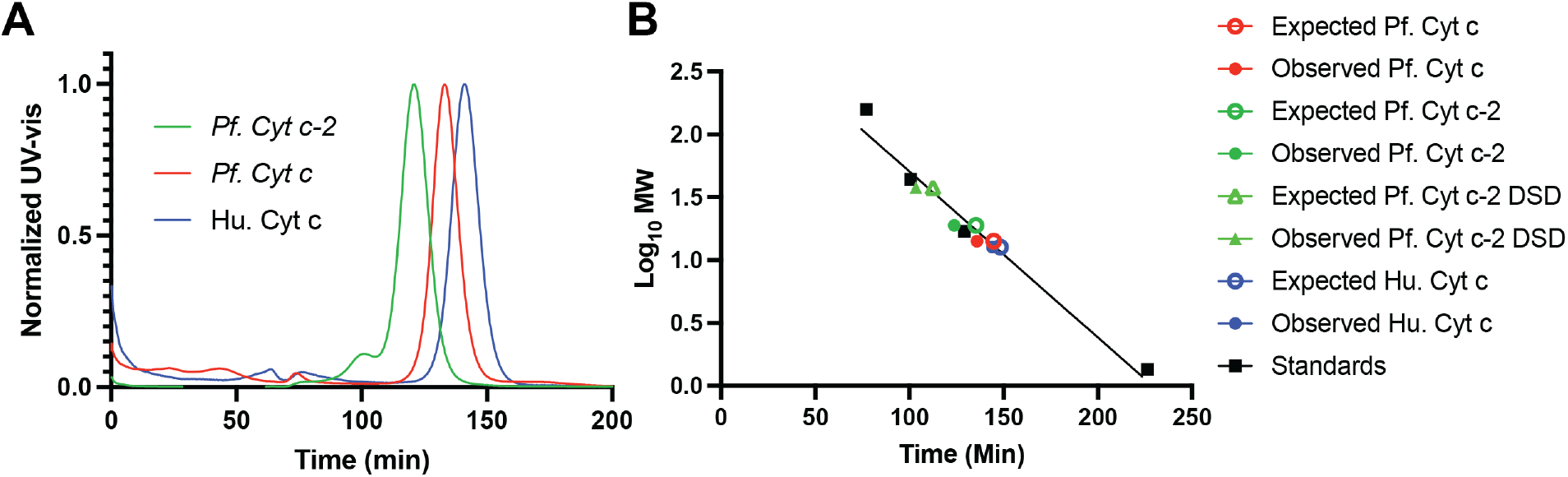
Size-exclusion chromatography of cyt *c* homologs. (A) Chromatograph of purified proteins run individually on an S-100 column. (B) Calibration curve for low MW versus retention time (RT) of globular proteins run on the S-100 column. This curve in black was used to interpolate the expected RT of the cytochrome c homologs based on MW and compared to the observed RT from panel (A), including the predicted RT of the cyt *c*-2 domain-swapped dimer (DSD).

**Figure S5.**
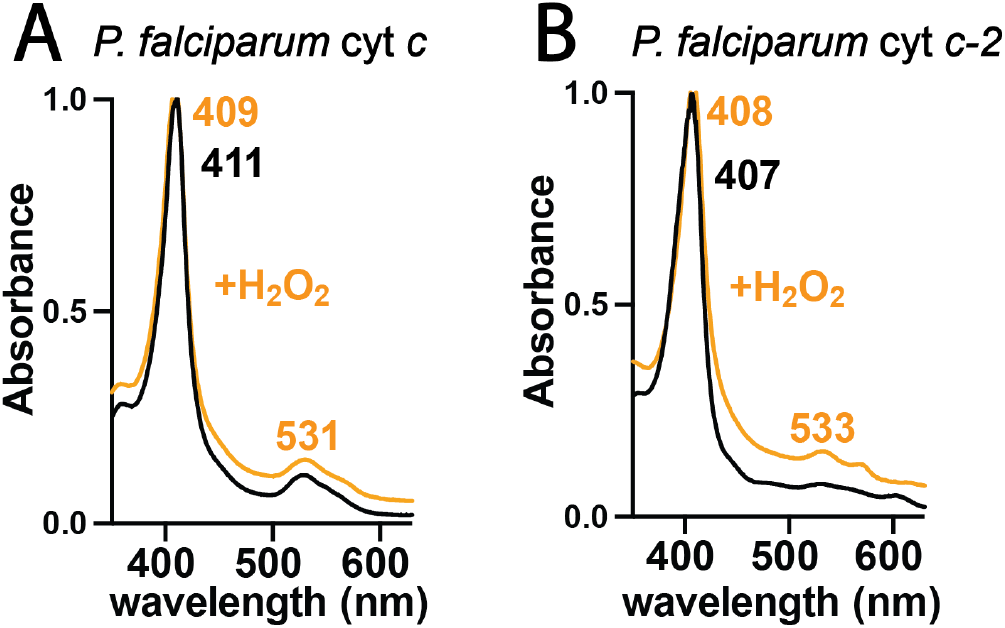
UV-vis absorbance spectrum of 10 μM cyt *c* (A) and cyt *c*-2 (B) oxidized with 1 mM ammonium persulfate (black) and then with 1 mM hydrogen peroxide (orange). Spectra were normalized to the Soret peak and offset vertically to avoid overlap.

**Figure S6.**
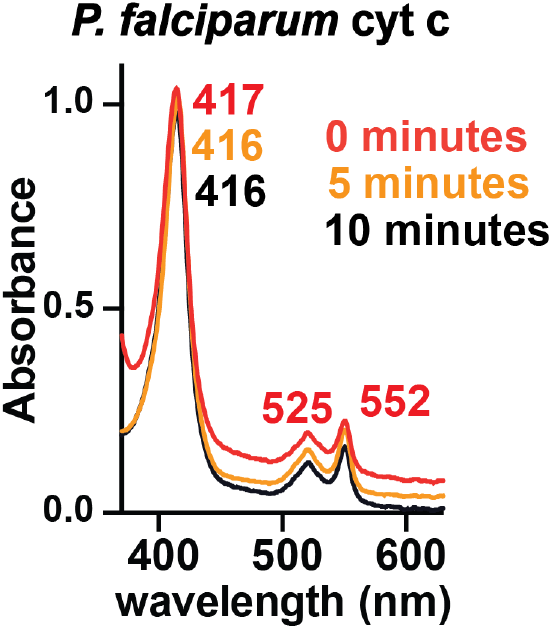
Time-dependent UV-vis absorbance spectra of parasite cyt *c* reduced by 1 mM sodium dithionite and exposed to air.

**Figure S7.**
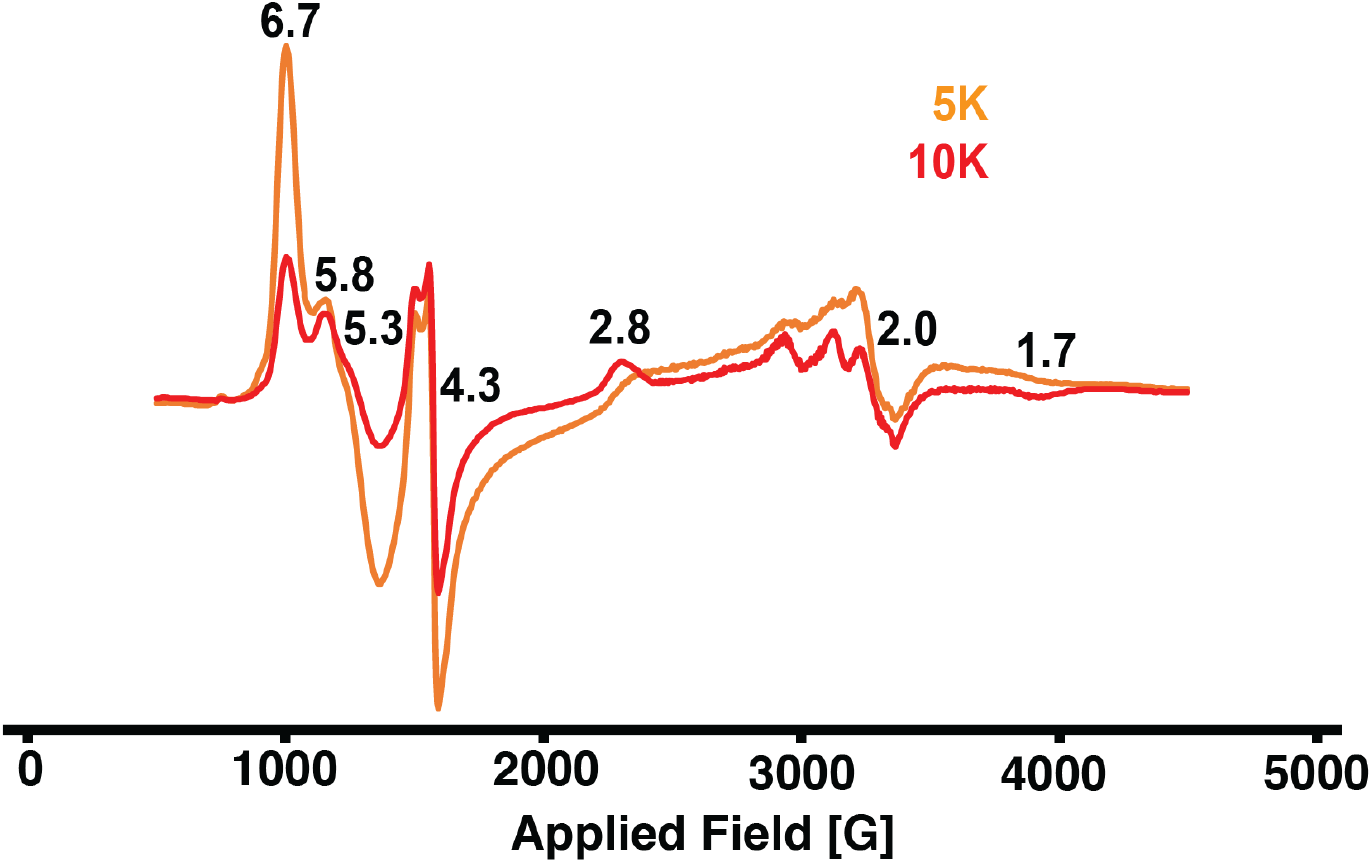
EPR spectra of cyt *c*-2 acquired at 5K or 10K and 1mW. EPR g values are indicated above major features.

**Figure S8.**
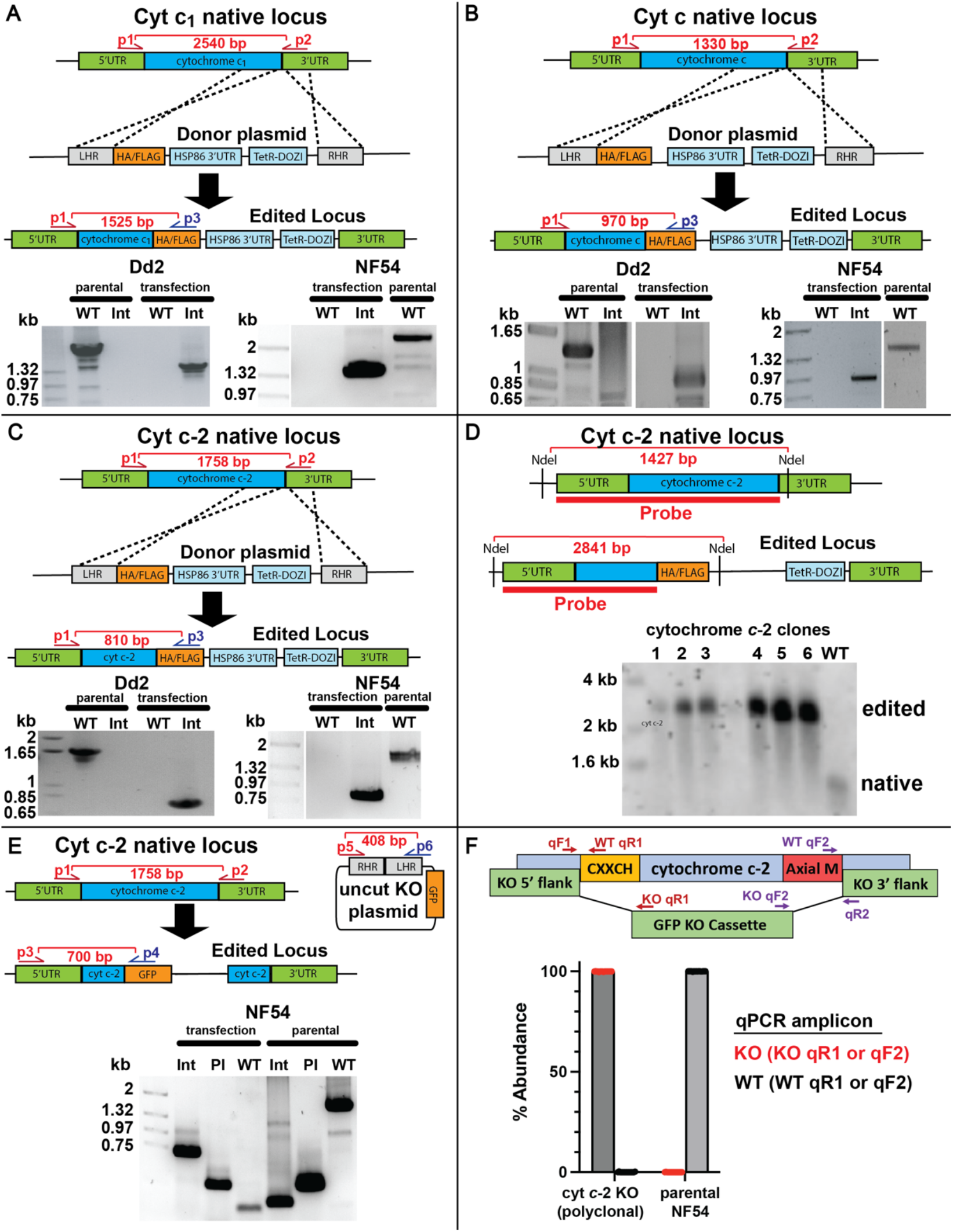
Confirmation of parasite genome editing by PCR and Southern blot. PCR analysis of genomic integration of the aptamer/TetR-DOZI system for polyclonal transfections of cyt *c*_1_ (A), cyt *c* (B), and cyt *c*-2 (C) in Dd2 and NF54 parasites. Approximate primer sites and expected amplicon sizes are indicated in scheme. For each analysis of Dd2 or NF54 parasites, samples were run on the same gel but images were cropped for clarity. (D) Southern blot analysis of genomic integration of the aptamer/TetR-DOZI system at the cyt *c*-2 locus in clonal Dd2 parasites. Approximate restriction sites, expected fragment sizes, and probe location are indicated in the scheme. PCR (E) and qPCR (F) analyses of genomic disruption of the cyt *c*-2 coding sequence in polyclonal NF54 parasites. The analysis (E) also contained a positive control reaction with added, uncut donor plasmid. Int = integrated, edited locus, WT = wildtype locus, Pl = uncut plasmid.

**Figure S9.**
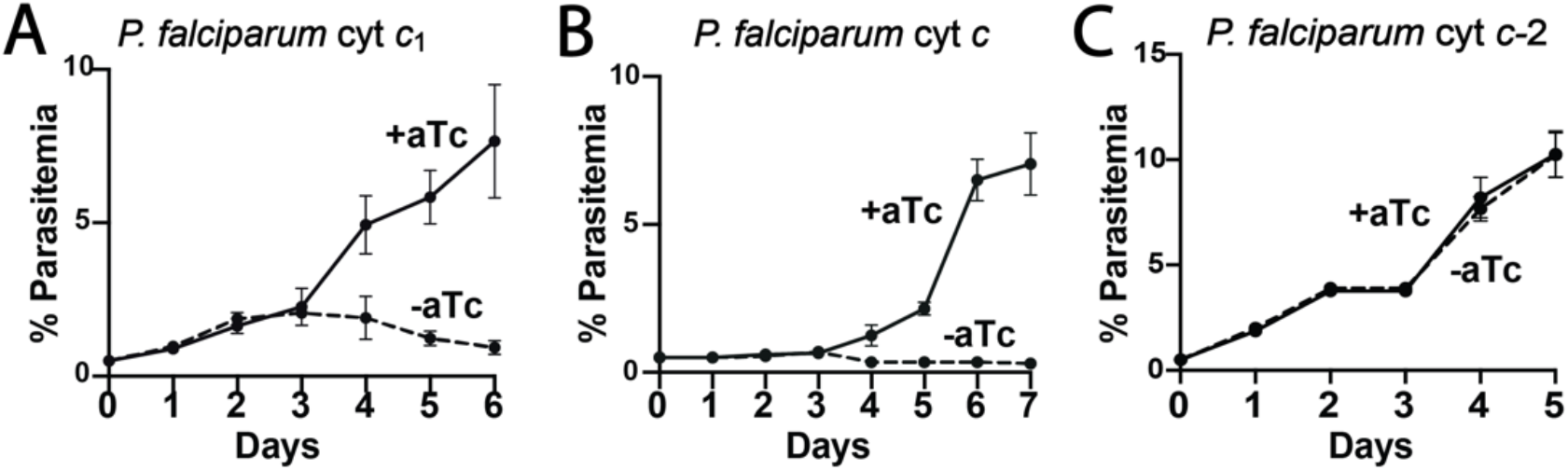
Functional tests of contributions by cyt *c*, *c*-2, and *c*_1_ to blood-stage parasite growth by NF54 parasites. Synchronous growth assays of NF54 parasites tagged with the aptamer/TetR-DOZI cassette for inducible knockdown of cyt *c*_1_ (A), cyt *c* (B), and cyt *c*-2 (C), in the presence or absence of 0.5 μM anhydrotetracycline (aTc).

**Figure S10.**
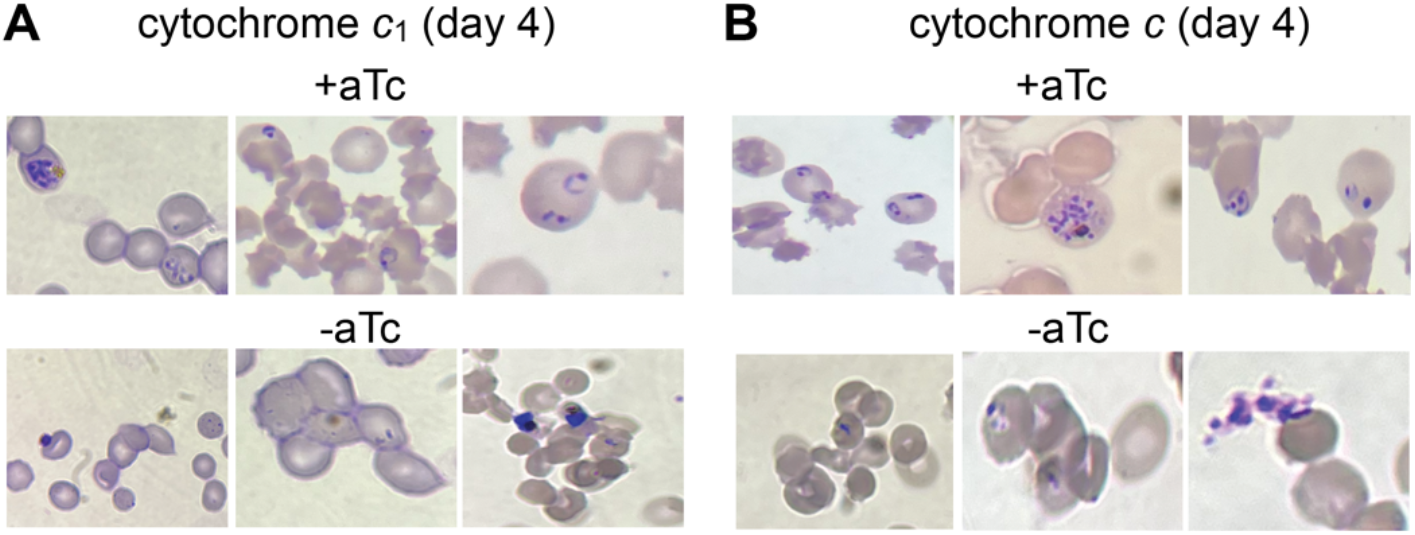
Blood-smear analysis of Dd2 parasites upon regulation of cyt *c*_1_ (A) and cyt *c* (B) expression after 4 days of growth ±aTc.

**Figure S11.**
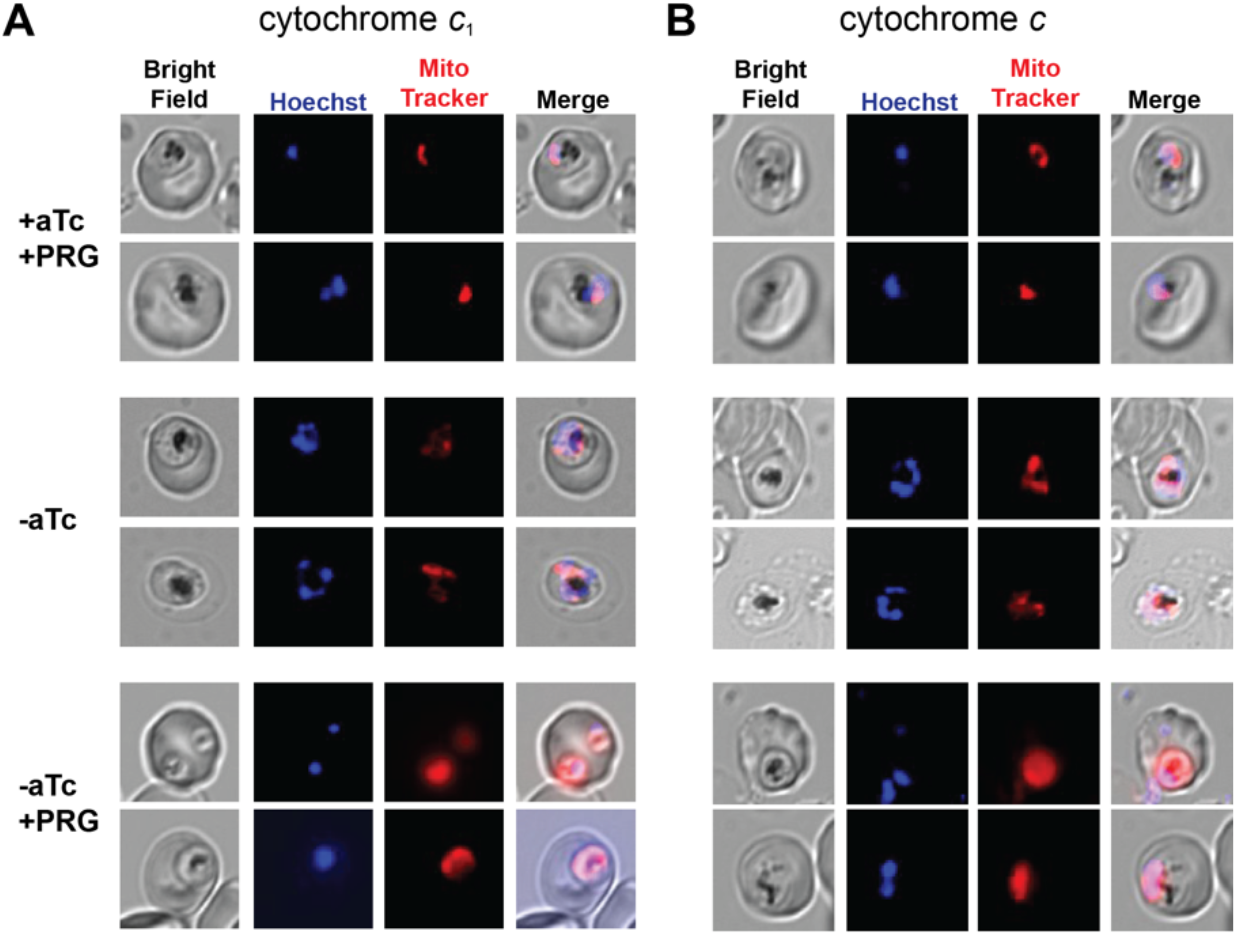
Additional live microscopy images of cyt *c*_1_ (A) and cyt *c* (B) aptamer/TetR-DOZI parasites cultured 3 days ±aTc and ± 1 μM proguanil and stained with Hoechst or MitoTracker Red (10 nM).

**Table S1.** Data collection and refinement statistics for determination of the cyt *c*-2 structure.

| Data collection and structure refinement statistics. |  |  |
| --- | --- | --- |
| Crystal Structure | Pf. Cyt *c*-2 DSD<br>(PDB: 7U2V) | Pf. cyt *c*-2 DSD<br>(PDB: 7TXE) |
| Beamline | SSRL 12-1 | SSRL 9-2 |
| Wavelength (Å) | 1.73816 | 0.97946 |
| Resolution range (Å) | 38.3 - 2.6 (2.6 - 2.6) | 27.2 - 2.3 (2.4 - 2.3) |
| Space group | P 1 21 1 | P 21 21 21 |
| Unit cell (Å) | 40.8, 54.6, 71.5 | 54.4, 72.0, 78.4 |
| Unit cell (°) | 90, 102.2, 90 | 90, 90, 90 |
| Total reflections | 100700 (2486) | 1451958 (144018) |
| Unique reflections | 8410 (334) | 14199 (1402) |
| Multiplicity | 12.0 (7.7) | 102.3 (102.9) |
| Completeness (%) | 83.8 (33.1) | 98.9 (99.3) |
| Mean I/sigma(I) | 20.5 (4.5) | 19.9 (2.3) |
| R-meas | 0.1783 (0.5167) | 0.4416 (4.778) |
| R-pim | 0.0503 (0.1849) | 0.0432 (0.4665) |
| Reflections used in refinement | 8509 (330) | 14076 (1393) |
| Reflections used for R-free | 850 (36) | 1401 (139) |
| R-work (%) | 19.00 (18.58) | 23.23 (31.32) |
| R-free (%) | 24.23 (22.06) | 28.16 (36.93) |
| RMS (bonds) | 0.004 | 0.005 |
| RMS (angles) | 0.49 | 0.6 |
| Ramachandran favored (%) | 98.6 | 95.74 |
| Ramachandran outliers (%) | 0 | 0 |
| Average B-factor (Å<sup>2</sup>) | 22.98 | 52.71 |

\*Statistics for the highest-resolution shell are shown in parentheses.

**Table S2.**
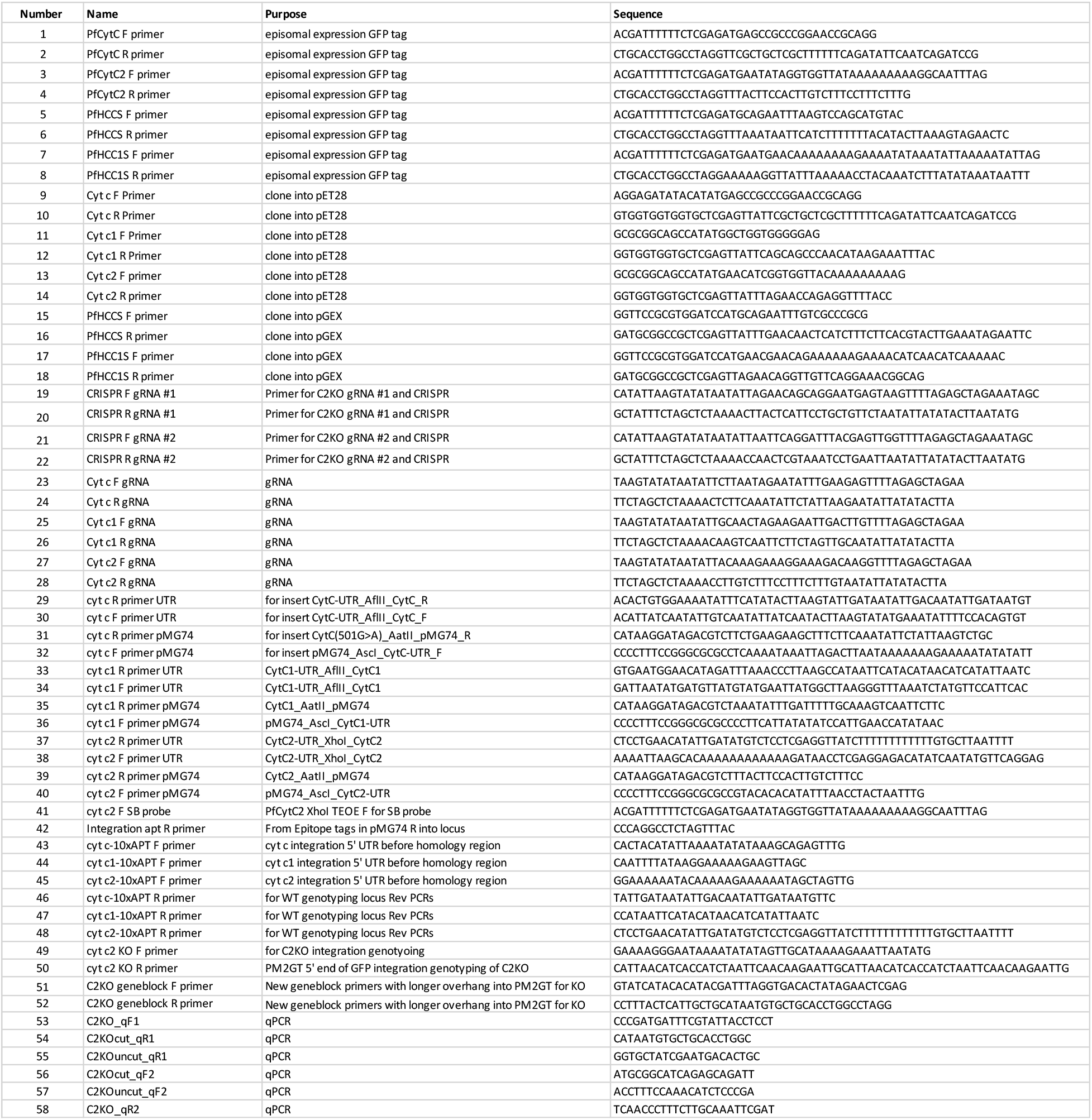
Table of PCR primers used for cloning, sequencing, and PCR analyses.

